# Germline Missense Variants in *CDC20* Result in Aberrant Mitotic Progression and Familial Cancer

**DOI:** 10.1101/2021.09.13.459529

**Authors:** Owen J. Chen, Ester Castellsagué, Mohamed Moustafa-Kamal, Javad Nadaf, Barbara Rivera, Somayyeh Fahiminiya, Yilin Wang, Isabelle Gamache, Caterina Pacifico, Lai Jiang, Jian Carrot-Zhang, Leora Witkowski, Albert M. Berghuis, Stefan Schoenberger, Dominik Schneider, Susanne Bens, Reiner Siebert, Colin J. R. Stewart, Ziguo Zhang, William C. Chao, Celia M.T. Greenwood, David Barford, Marc Tischkowitz, Jacek Majewski, William D. Foulkes, Jose G. Teodoro

## Abstract

CDC20 is a co-activator of the anaphase promoting complex/cyclosome (APC/C) and is essential for mitotic progression. APC/C^CDC20^ is inhibited by the spindle assembly checkpoint (SAC), which prevents premature separation of sister chromatids and aneuploidy in daughter cells. Although overexpression of *CDC20* is common in many cancers, oncogenic mutations have never been identified in humans. Using whole exome sequencing, we identified heterozygous missense *CDC20* variants (L151R and N331K) that segregate with cancer in two families. Characterization of these mutants showed they retain APC/C activation activity but show reduced binding to BUBR1, a component of the SAC. Expression of L151R and N331K promoted mitotic slippage in HeLa cells and primary skin fibroblasts derived from carriers. CRISPR/Cas9 was used to generate mice carrying N331K. Homozygous mice carrying N331K were non-viable, however, heterozygotes displayed accelerated oncogenicity in Myc-driven cancers. These findings highlight an unappreciated role for *CDC20* variants as tumor promoting genes in humans.

## INTRODUCTION

CDC20 (cell division cycle 20 homolog) is a co-activator of the anaphase promoting complex/cyclosome (APC/C).^1^ The APC/C is a large multi-subunit E3 ubiquitin ligase that is critical for cell cycle regulation.^2^ During mitosis, CDC20 binds and activates the APC/C, which ubiquitinates mitotic substrates cyclin B and securin in order for sister chromatids to separate and anaphase to proceed.^3,4^ Mitotic progression is regulated by the spindle assembly checkpoint (SAC), also known as the mitotic checkpoint.^3,4^ The SAC is triggered when cells in mitosis are not ready to proceed to anaphase, such as when kinetochores of chromosomes are not properly attached to the spindle fibers during prometaphase.^3,4^ Kinetochores that lack spindle attachment provide a negative signal to the APC/C^CDC20^ complex through the mitotic checkpoint complex (MCC), formed by the proteins MAD2, BUBR1 and BUB3, along with CDC20.^5,6^ The MCC binds APC/C^CDC20^ directly and inhibits ubiquitination activity, resulting in a delay in anaphase until all chromosomes are properly attached to the spindles.^7,8^ The timing of sister chromatid separation must be perfect in order to maintain proper ploidy.^9^ All eukaryotic cells have this fail-safe mechanism to prevent aberrant chromosomal segregation.^10,11^ Dysregulation of CDC20 and APC/C activity can lead to problems in chromosome segregation, mitotic progression, and cell viability.^12,13^ These aberrant events have many pathological implications, including aneuploidy and tumorigenesis.^12^ However, mutations in genes coding for mitotic checkpoint proteins such as CDC20 are often embryonic lethal as they are essential to life.^12,14,15^ Thus, such mutations are rare in human disease,^12,16^ and the clinical significance of mutations in *CDC20* have been investigated in only a handful of studies in an animal model or human context.^17–23^ Although *CDC20* is overexpressed in the majority of human cancers,^16,24–30^ the involvement of *CDC20* mutations in cancer has never been established in humans.

In the current study, we originally sought to identify the underlying genetic causes of familial malignant ovarian germ cell tumors (mOGCTs). We identified 8 families with histories of mOGCT.^31^ We were able to obtain biological material from 5 of these families and performed whole exome sequencing (WES) on blood DNA to look for likely pathogenic or pathogenic variants that segregated in one or more families. Using this approach, two distinct missense mutations in the *CDC20* gene were identified in all affected individuals of 2 families. Since aneuploidy and chromosome instability are implicated in mitotic checkpoint defects and across different cancers,^11^ we sought to investigate the role of *CDC20* mutations in cancer patients on cell cycle dysregulation and tumorigenesis. In parallel, we performed WES on germ cell tumors (GCTs) of 85 other individuals with sporadic cases to look for other *CDC20* mutations. Here, we report our molecular and *in vivo* findings from our study of two germline *CDC20* mutations in patients from two mOGCT families.

## RESULTS

### Identification of *CDC20* variants in two families with histories of cancer

Saliva or blood samples were collected from five out of eight families previously reported^31^ to show mOGCT aggregation and WES analysis was performed on the germline DNA of available affected members (**Figures 1A and B, S1, S2, S3 and Table S1A**). Two distinct *CDC20* missense variants were identified in all affected members of two (‘fmOGCT4’ and ‘fmOGCT7’) of the five families studied, one variant in each family (**Table S1B and C**). No other variants in this gene that have met our filtering criteria were found in any of the other three analyzed families (**Table S1D, E and F**). Given that the probability of finding 2 out of 5 families with very rare (allele frequency ≤ 1×10^−5^) segregating variants in the same gene is low (p = 0.041) (**Table S2**), we selected the two *CDC20* variants for further analyses. Family fmOGCT4^32^ harbors the chr1:43825664T>G (hg19); c.452T>G (NM_001255.2); p.L151R variant (referred to as L151R onwards), which co-segregates with the disease in all affected women (**Figure 1A**). Only one male carries the mutation and remains unaffected, most likely due to incomplete penetrance or sex-related factors. Affected women in this family also presented with non-mOGCT cancers. In family fmOGCT7,^33^ two affected siblings inherited the chr1:43826548C>G (hg19); c.993C>G (NM_001255.2); p.N331K variant (referred to as N331K onwards) from their mother who presented with endometriotic cysts and multinodular goiter. Of note, the family presented with a history of goiter, but whether *CDC20* is related to that phenotype remains uncertain (**Figure 1B**). Both variants from both families were heterozygous, expressed at the RNA level, and no loss of the wild-type (WT) allele or second hit in *CDC20* was found in the tumors analyzed, thus suggesting a dominant mode of inheritance (**Figure 1A and B**). In addition, both variants were predicted to be pathogenic by several *in silico* models (see **Figure 1C** legend).

**Figure 1:**
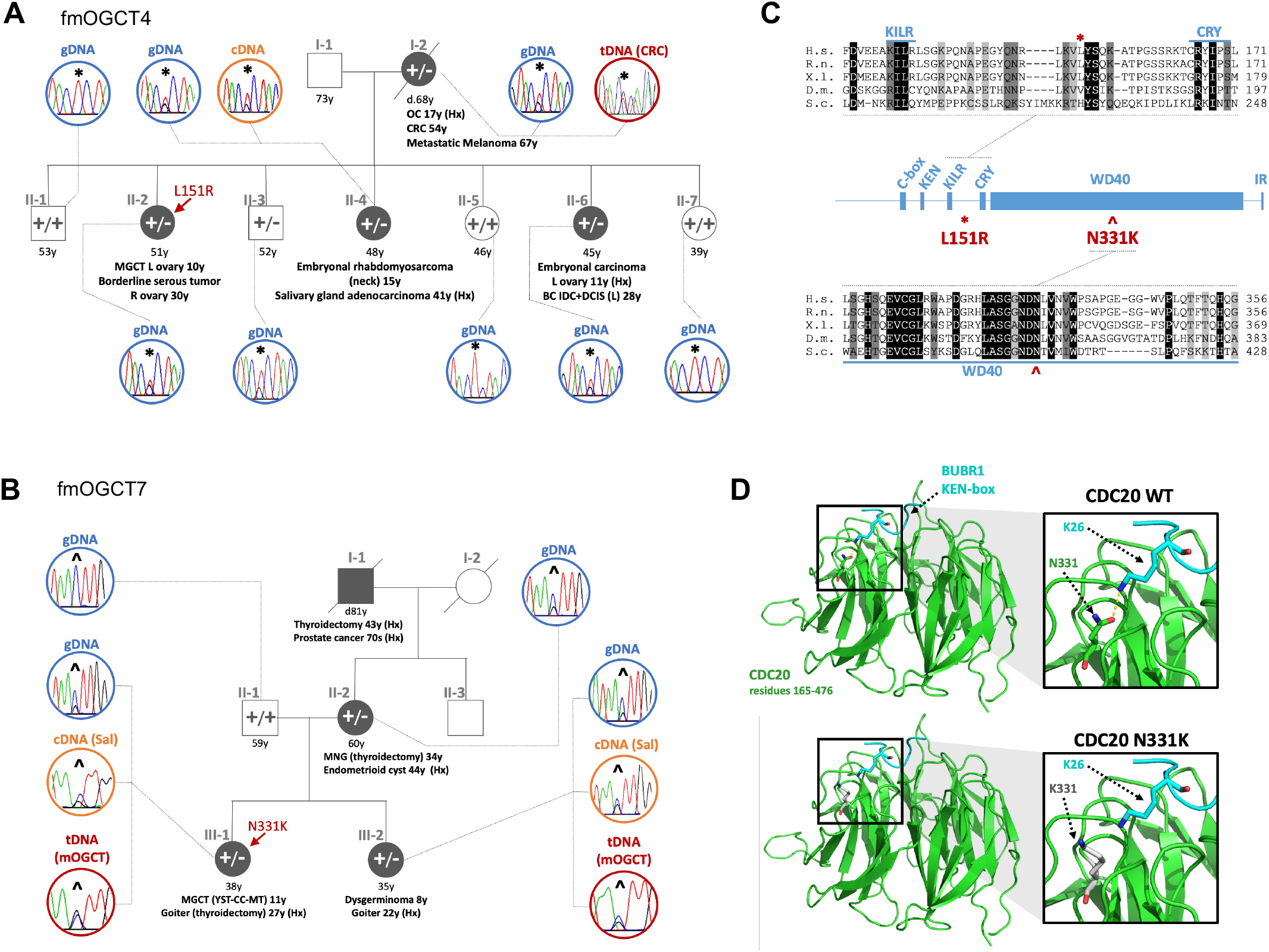
Pedigrees of the *CDC20* mutation carriers and locations of the L151R and N331K mutations. **(A and B)** In panel **A**, fmOGCT4 carries the c.452T>G, p.L151R mutation (*). In panel **B**, fmOGCT7 carries the c.993C>G, p.N331K mutation (^). In both families, the mutation correlates with the disease, and is expressed and shows no loss of heterozygosity in the tumors analyzed. Probands are identified with an arrow. Affected individuals are filled in grey. Mutated and WT *CDC20* alleles are shown as minus and plus signs, respectively. Age at last follow-up or age at death appears in years below for each individual. Age at diagnosis is shown after disease description below affected individuals. Chromatograms from Sanger sequencing of different tissue types also appear for several individuals. OC: ovarian cancer; CRC: colorectal cancer; GCT: germ cell tumor; R: right; L: left; BC: breast cancer, IDC: invasive ductal carcinoma; DCIS: ductal carcinoma *in situ*; MNG: multinodular goiter; MGCT: mixed GCT; YST: yolk sac tumor; CC: choriocarcinoma; Hx: family history, or otherwise confirmed by clinical records; gDNA: genomic DNA; cDNA: complementary DNA obtained from RNA; tDNA: DNA from tumoral tissue; Sal: saliva. **(C)** Schematic representation of the domains and motifs of human CDC20: C-box and IR tail are important for association with APC/C; KEN and CRY sequences are degron recognition sites essential for CDC20 degradation; KILR sequence is part of the MAD2-interacting motif; seven WD40 repeats are important for mediating interaction with other proteins such as BUBR1. Multiple sequence alignment from different organisms of the CDC20 residues surrounding the two familial mutations were performed using CLUSTAL O (1.2.4). The black background indicates positions with fully conserved residues, dark grey indicates conservation of residues of strongly similar properties, and light grey indicates conservation of residues of weakly similar properties. The L151R and N331K variants were predicted to be pathogenic by several *in silico* models: Combined Annotation Dependent Depletion (CADD) scores for L151R and N331K are 26 and 25, respectively; Revels scores for L151R and N331K are 0.47 and 0.15, respectively. A CADD score ≥ 20 indicates a deleterious variant. Revel scores range from 0 to 1, and more damaging variants have higher scores. H.s.: *Homo sapiens*; R.n.: *Rattus norvegicus*; X.l.: *Xenopus*; D.m.: *Drosophila melanogaster*; S.c.: *Saccharomyces cerevisiae*. **(D)** Computational structural modelling comparing the theoretical protein structures of WT CDC20 and the N331K variant (green), and their interactions with the KEN1-box of BUBR1 (cyan). The interactions between residues N331 (in CDC20) and K26 (in BUBR1) are highlighted in the insets. The hydrogen bond formed between WT CDC20 and BUBR1 is illustrated by the dashed yellow line. The CDC20 N331K mutation is predicted to prevent this hydrogen bond interaction with BUBR1.

In view of these results, we also screened 85 sporadic GCTs for *CDC20* mutations and found a predicted deleterious mutation in one tumor belonging to a male patient (c.947G>A, p.R316H). Germline DNA was not available in this case. However, there was no evidence that this variant has any functional implications (data not shown).

WES was also performed on available tumoral material: fresh frozen tissue from fmOGCT7: III-1(mixed GCT) and fmOGCT7: III-2 (Dysgerminoma, DG). As putative driver mutations, we found a variant in *AKT2* in the mixed GCT and a variant in *KIT* in the DG (**Table S3**). Genome-wide tumoral interrogation was done using the OncoScan microarray assay on FFPE (formalin-fixed paraffin-embedded) samples from these two tumors. Both showed several gains and/or losses mostly affecting whole chromosomes or chromosome arms, resulting in considerable aneuploidy overall and consistent with *CDC20* disruption (**Figure 2A**). The copy number pattern of fmOGCT7 showed shared gains involving chr7, 8, 12 and 20 and a partial deletion region on p-arm of chr3 in both sisters (**Figure 2B**). Aneuploidy was also detected with the FISH analysis performed on available tumoral slides, which showed a significant gain of chromosomes in the two mOGCTs analyzed (**Table S4**). Furthermore, blood and skin cell samples were taken from the two sisters (III-1 and III-2) in fmOGCT7 and the karyotypes were analyzed in 50 cells for each. Chromosomal analyses revealed single aneuploid cells as well as subclones with a t(3;13) and trisomy X in fibroblast and lymphocyte cultures, respectively (data not shown), from III-1 (**Figure 1B**).

**Figure 2:**
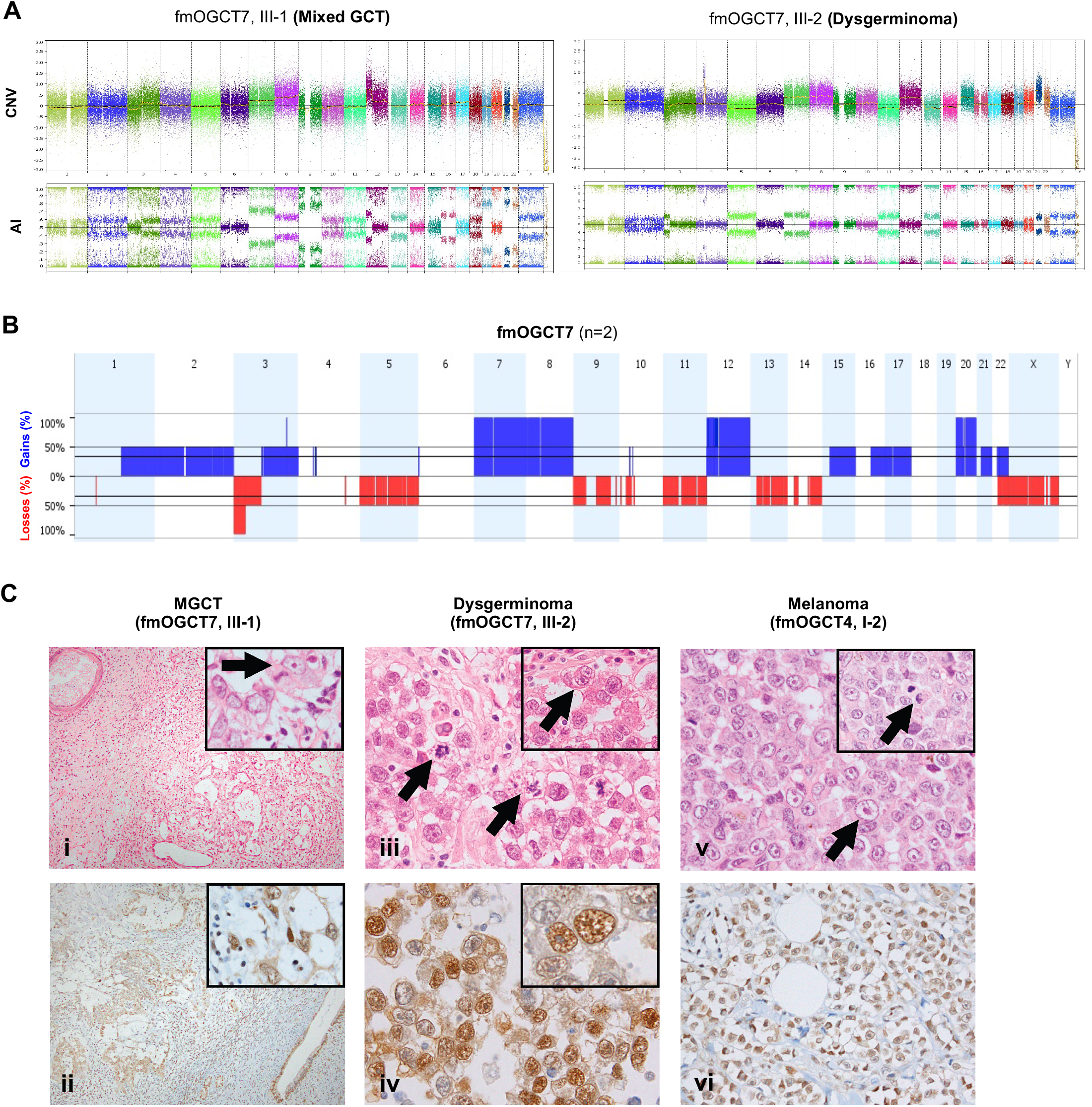
Tumoral tissues from mOGCT patients show chromosomal abnormalities. **(A)** OncoScan analysis on tumors. Copy number changes (CNV) and allelic imbalance (AI) are shown for the two affected sisters: fmOGCT7_III-1 and fmOGCT7_III-2. The y-axes of the CNV and AI plots indicate log2 ratios and B-allele frequency (BAF) ranging from 0 to 1 (with expected BAF value of 0.5 for heterozygous loci), respectively. Both samples show CNV on several chromosomes, which are also reflected in the AI profiles. **(B)** Copy number profiles for both samples (affected siblings of fmOGCT7). Shared copy number gains (blue) can be observed on chromosomes 7, 8, 12, 20, as well as a loss (red) on chromosome 3. **(C)** H&E staining and CDC20 immunohistochemistry on patient tumor sections. **(i & ii)**: Malignant mixed germ cell tumor (fmOGCT7, III-1). **(i)** shows histologically mature elements (squamous lined cyst, upper left) and yolk sac tumor (YST) (lower right). The inset shows cytologic detail of the YST. Vesicular nuclei, prominent eosinophilic nucleoli, and peri-nucleolar clearing are shown with the arrow. **(ii)** is CDC20 immunohistochemistry showing nuclear and cytoplasmic staining in the YST (left) and in an adjacent benign gland (right). The inset shows granular nuclear immunoreactivity in the YST. **(iii & iv)**: Dysgerminoma (fmOGCT7, III-2). In **(iii)**, numerous mitotic figures are present including abnormal partial ring forms (left arrow) and asymmetric divisions (right arrow). The inset shows cytologic detail including cells with prominent nucleoli and peri-nucleolar clearing (arrow). In **(iv)**, immunohistochemistry shows nuclear CDC20 expression in most tumor cells together with weaker cytoplasmic staining. The inset demonstrates the granular nuclear staining pattern. **(v & vi)**: Metastatic melanoma (fmOGCT4, I-2). In **(v)**, many tumor cells show large vesicular nuclei, prominent nucleoli, and peri-nucleolar clearing (arrow). The inset shows an abnormal, asymmetric mitotic figure (arrow). **(vi)** shows CDC20 immunohistochemistry of melanoma within subcutaneous fat. Most tumor cells show moderate nuclear and cytoplasmic staining.

Immunohistochemistry (IHC) was performed to visualize CDC20 on mOGCTs derived from III-1 and III-2 from fmOGCT7, the metastatic melanoma in I-2 from fmOGCT4, and a tissue microarray (TMA) of a series of sporadic mOGCTs. Many of the mitotic figures in the *CDC20*-mututed tumors, particularly the mOGCTs, were morphologically abnormal with asymmetric divisions and occasional partial or complete ring forms, which is consistent with dysregulated CDC20 function (**Figure 2C**). Some tumor cells also demonstrated distinctive vesicular nuclei with prominent eosinophilic nucleoli and peri-nucleolar chromatin clearing. More than 50% of the cells in the familial tumors demonstrated nuclear and cytoplasmic CDC20 immunoreactivity and the mOGCTs showed an unusual granular/reticular nuclear staining pattern compared to the more diffuse immunoreactivity seen in most other sporadic tumors (**Figure 2C and Table S5**). However, there were no significant differences in overall staining extent or intensity between the familial cases and the sporadic mOGCT cases evaluated in the TMA (**Table S5**). Taken together, these findings suggest that *CDC20* mutations may play a role in the tumorigenesis in the aforementioned familial cancers by disrupting the normal function and regulation of CDC20 in mitosis.

### CDC20 variants compromise the spindle assembly checkpoint

Although the L151 residue is close to the MAD2-binding domain (**Figure 1C**), L151R could not be modelled structurally due to its location in a disorganized part of the protein. In contrast, the N331 residue is in the highly conserved KEN-box binding site and forms a direct hydrogen bond with the K26 residue of the KEN1 motif of BUBR1.^5^ Computational structural analysis showed that mutating the N331 residue to a lysine (K) removes this hydrogen bond interaction, creating an electrostatic repulsion that could potentially weaken the affinity between CDC20 and BUBR1 (**Figure 1D)**.

In order to determine if the identified CDC20 variants affect the SAC, HeLa cells were transfected with FLAG-tagged CDC20 constructs expressing WT CDC20 or other variants: L151R and N331K, present in the familial cases, and R132A. R132A is a well characterized mutation with defective binding to MAD2 which serves as a positive control for SAC disruption.^8^ Transfected cells were synchronized with thymidine (Thy), released, and arrested in mitosis with nocodazole (NOC). The mitotic status of cells was monitored by immunoblot detecting the phosphorylated forms of BUBR1 and APC3 as mitotic markers. Transfected cells expressing each of the mutant CDC20 constructs displayed increased levels of the unphosphorylated forms of BUBR1 and APC3 compared to WT (**Figure 3A**, compare lane 4 to lanes 5, 6, and 7). In addition, flow cytometry performed on the Thy-NOC synchronized transfected cells, showed that cells expressing the CDC20 mutants displayed a greater proportion of cells with tetraploidy, suggesting that the mutant CDC20 expressing cells were more refractive to engagement of the SAC by NOC (**Figure 3B and C**). In order to better characterize perturbations in the cell cycle, live cell, time-lapse microscopy, was used to image HeLa cells transfected with bicistronic vectors expressing either WT or mutant CDC20 (see Methods) for 40 h after Thy-NOC synchronization. **Figure 3D and E** shows that cells transfected with mutant CDC20 underwent significantly more mitotic slippage and less mitotic cell death compared to cells transfected with WT. The elevated levels of slippage observed in cells expressing the L151R and N331K mutations suggested that the SAC was attenuated. To test this possibility, immunoprecipitations (IPs) were performed on FLAG-tagged CDC20 from transfected HeLa mitotic cell extracts followed by immunoblot to detect components of the SAC. We observed that both the N331K and L151R mutants had decreased affinity to BUBR1 during mitosis (**Figure 3F**, lanes 3 and 4; **Figure S4A**, lanes 3 and 4). As expected, the R132A CDC20 mutant had a reduced interaction with MAD2 (**Figure 3F**, lane 5). These data suggest that the L151R and N331K mutations disrupt normal mitotic progression by compromising interaction with the SAC.

**Figure 3:**
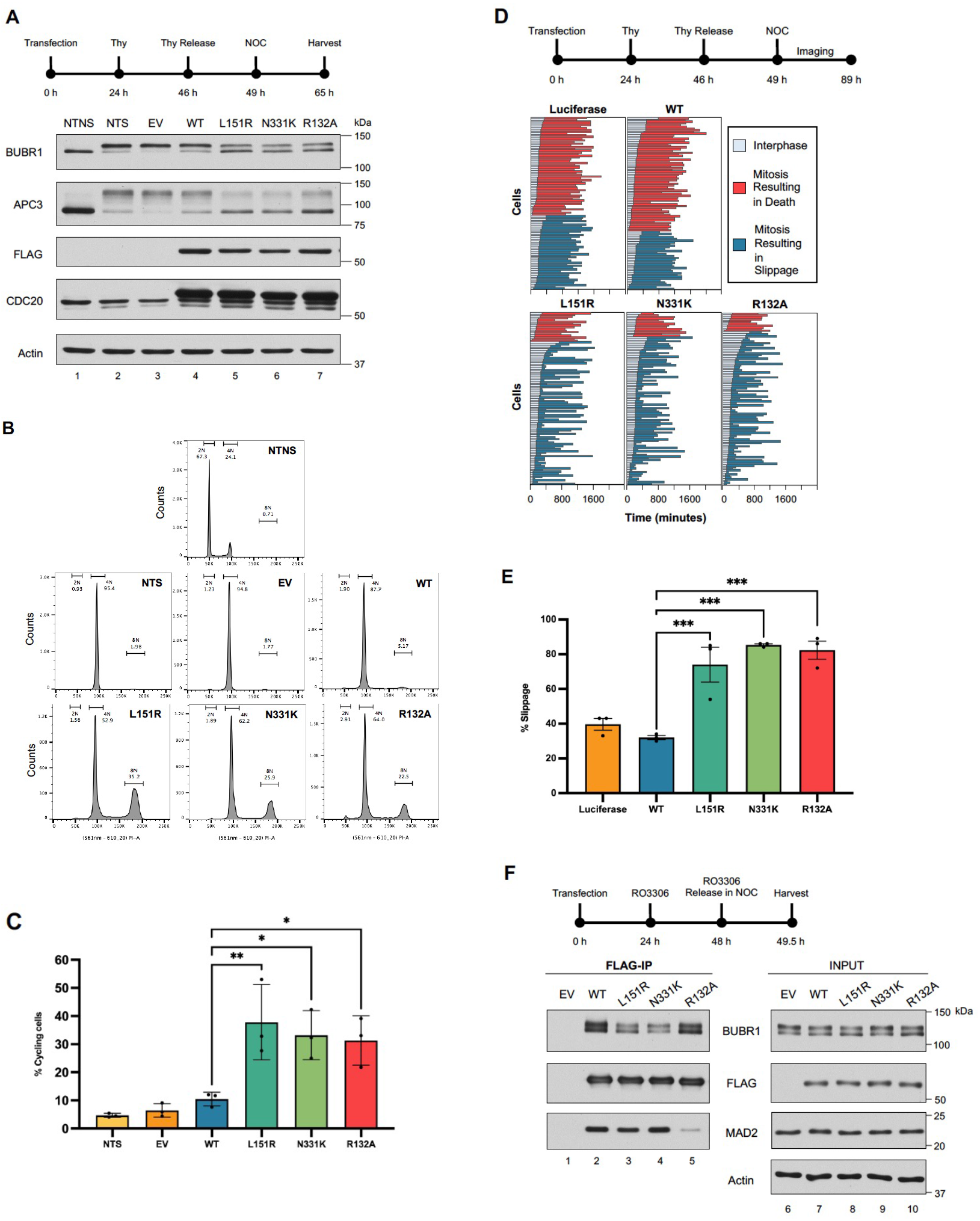
The L151R and N331K CDC20 variants from mOGCT patients compromise the SAC in HeLa cells. **(A)** Expression blots of extracts of cells transfected with FLAG-tagged WT CDC20 or mutant constructs and synchronized with thymidine (Thy) and nocodazole (NOC). A visual representation of the experimental procedure is included above. The phosphorylated forms of BUBR1 and APC3 are used as mitotic markers. NTNS: non-transfected, non-synchronized; NTS: non-transfected, synchronized; EV: empty vector. **(B)** Flow cytometry analysis of transfected, synchronized cells (same treatment protocol as in **B**) stained with propidium iodide for DNA content. The 8N peaks indicate non-mitotic, polyploid (>4N) cells. **(C)** Quantification of the proportion of non-mitotic, tetraploid cells from the flow cytometry analysis in **B**, representing 3 independent experiments. An ordinary one-way ANOVA with Dunnett’s multiple comparisons test was used (n = 3; *: p < 0.05; **: p < 0.01). Error bars: SD. **(D)** Visual representation of the fates of individual cells transfected with bicistronic constructs expressing the different CDC20 mutants and GFP observed using real-time microscopy for 40 h after the addition of nocodazole to synchronized cells. A visual timeline of the experimental procedure of this mitotic slippage assay is included above. Images were taken every 10 min, and each analyzed GFP-positive cell was noted for the frame at which it entered mitosis, and when it underwent apoptosis during mitosis or slipped out of mitosis. Cells that died or slipped were not analyzed further. The colour scheme of the cell fates is as follows: grey, interphase; red, mitosis resulting in death; blue, mitosis resulting in mitotic slippage. The frames were converted to and shown in minutes (x-axis). N = 100 cells per construct (y-axis). These results are representative of 3 independent time-lapse experiments. **(E)** Quantification of the proportion of cells that underwent mitotic slippage across the different CDC20 mutants from the time-lapse experiment in panel **D**. An ordinary one-way ANOVA with Dunnett’s multiple comparisons test was used (n = 3; ***: p < 0.001). Error bars: SEM. **(F)** Immunoblots of different mitotic proteins from FLAG co-immunoprecipitations (co-IPs) on mitotic extracts from cells transfected with FLAG-tagged CDC20 WT or mutant constructs. The visual representation of the experimental procedure is included above. EV: empty vector.

### Familial CDC20 variants retain APC/C co-activator activity, which is required to promote mitotic slippage

The L151R and N331K mutants were able to attenuate the SAC in HeLa cells and promote slippage, suggesting that the mutant CDC20 retained activity as an APC/C co-activator. To test this possibility, we reconstituted *Cdc20* conditional knockout (KO) mouse embryonic fibroblasts (MEFs) with the bicistronic CDC20 constructs as mentioned above. The *Cdc20* KO conditional MEFS are derived from mice in which exon 2 of the *Cdc20* gene is flanked by LoxP sequences and knock-out is induced by addition of 4-hydroxytamoxifen (4-OHT) to stimulate expression of Cre recombinase.^15^ As shown in **Figure 4A**, approximately 90% of cells become rounded and arrested in mitosis following 48 h of 4-OHT induction. MEFs were electroporated with WT or mutant bicistronic CDC20 constructs and analyzed by real-time microscopy as shown schematically in **Figure 4B. Figure 4C and D** show that reintroduction of WT CDC20 into CDC20 KO cells rescued mitotic arrest in approximately 50% of transfected cells. Reconstitution with the CDC20 point mutants L151R, N331K and R132A were all able to rescue mitotic arrest comparable to WT. In contrast, the mutant R499E, in which the IR tail of CDC20 is mutated, was completely defective for rescue. The IR amino acid motif is present on the C-termini of both APC/C co-activators, CDC20 and CDH1, and is essential for binding and activation of the APC/C.^34^ Immunoblotting was performed to confirm that the mutants expressed at similar levels (**Figure S4B**).

**Figure 4:**
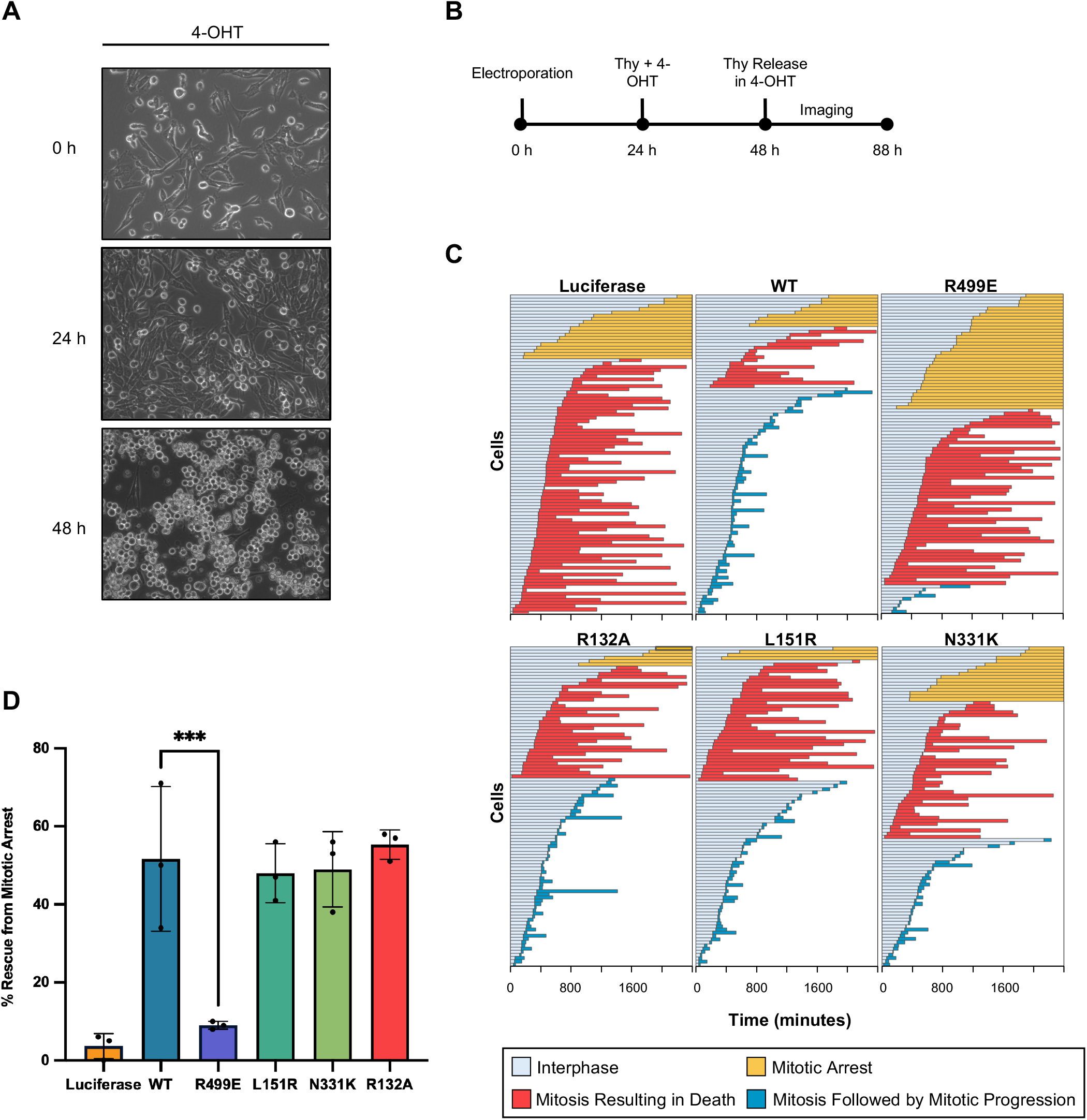
Familial *CDC20* variants retain APC/C co-activator activity. **(A)** Representative images confirming that the majority of *Cdc20* conditional knockout (KO) mouse embryonic fibroblasts (MEFs) treated with 4-OHT (4-hydroxytamoxifen) undergo mitotic arrest by 48 h. **(B)** Visual timeline of the experimental procedure of the mitotic rescue assay performed in *Cdc20* KO MEFs. Bicistronic constructs expressing the different CDC20 mutants with GFP were electroporated into the MEFs. **(C)** Visual representations of the fates of individual cells electroporated with bicistronic constructs expressing the different mutant CDC20 mutants with GFP observed using real-time microscopy for 40 h after thymidine release in 4-OHT. Images were taken every 10 min, and each analyzed GFP-positive cell was noted for the frame at which it entered mitosis, and when it underwent apoptosis during mitosis or exited mitosis. Cells that underwent mitotic progression were not followed further within the 40 h. Cells that stayed arrested in mitosis by the end of the 40 h were labelled as “mitotic arrest”. The colour scheme of the different cell fates is as follows: grey, interphase; orange, continued mitotic arrest; red, mitosis resulting in death; blue, mitosis resulting in mitotic progression (either cell division or mitotic slippage). The frames were converted to and shown in minutes (x-axis). N = 100 cells per construct (y-axis). These results are representative of 3 independent time-lapse experiments. **(D)** Comparison of the differences in percent rescue of GFP-positive cells from mitotic arrest for the different CDC20 mutants in the real-time microscopy experiment in **C**. An ordinary one-way ANOVA with Dunnett’s multiple comparisons test was used (n = 3; ***: p < 0.001). Error bars: SD.

To determine if APC/C binding activity is required for induction of mitotic slippage induced by the familial CDC20 variants, the L151R, N331K or R132A variants were combined with the R499E point mutation (**Figure 5A**). IPs were performed in mitotic cell extracts to confirm the effects of the mutations. MAD2 binding was decreased in the mutants that include R132A (**Figure 5B**, lanes 3, 4, and 7), and APC/C interaction was reduced in mutants containing the R499E as previously described^34^ (**Figure 5B**, lanes 5-7). All double mutants containing the L151R and N331K mutations had decreased affinity to BUBR1, as well as impaired interaction with BUB3 (**Figure 5B**, lanes 3-6). All double mutants expressed equally well (**Figure S4C**). **Figure 5C** shows that whereas the individual L151R, N331K or R132A mutants were able to significantly increase the level of mitotic slippage, combination of these mutations with the R499E IR tail mutation completely reversed this effect. In these experiments, slippage was also induced by the R499E mutant and notably, this effect was abrogated by combination with L151R, N331K or R132A, confirming that interaction with the SAC is essential for slippage induced by the IR tail mutant (**Figure 5B**). In addition to elevated slippage, time-lapse microscopy in HeLa cells also revealed that R132A, L151R and N331K had significantly shorter mitotic transit times in cells that underwent slippage in the presence of NOC (**Figure 5D**). The short mitotic transit times of the mutants were restored to WT levels when combined with the R499E mutation. Taken together, these data show that the CDC20 variants L151R and N331K compromise the mitotic checkpoint via impaired interaction with BUBR1 and that interaction and activation of the APC/C via the IR tail is required for this effect (**Figure 5E**).

**Figure 5:**
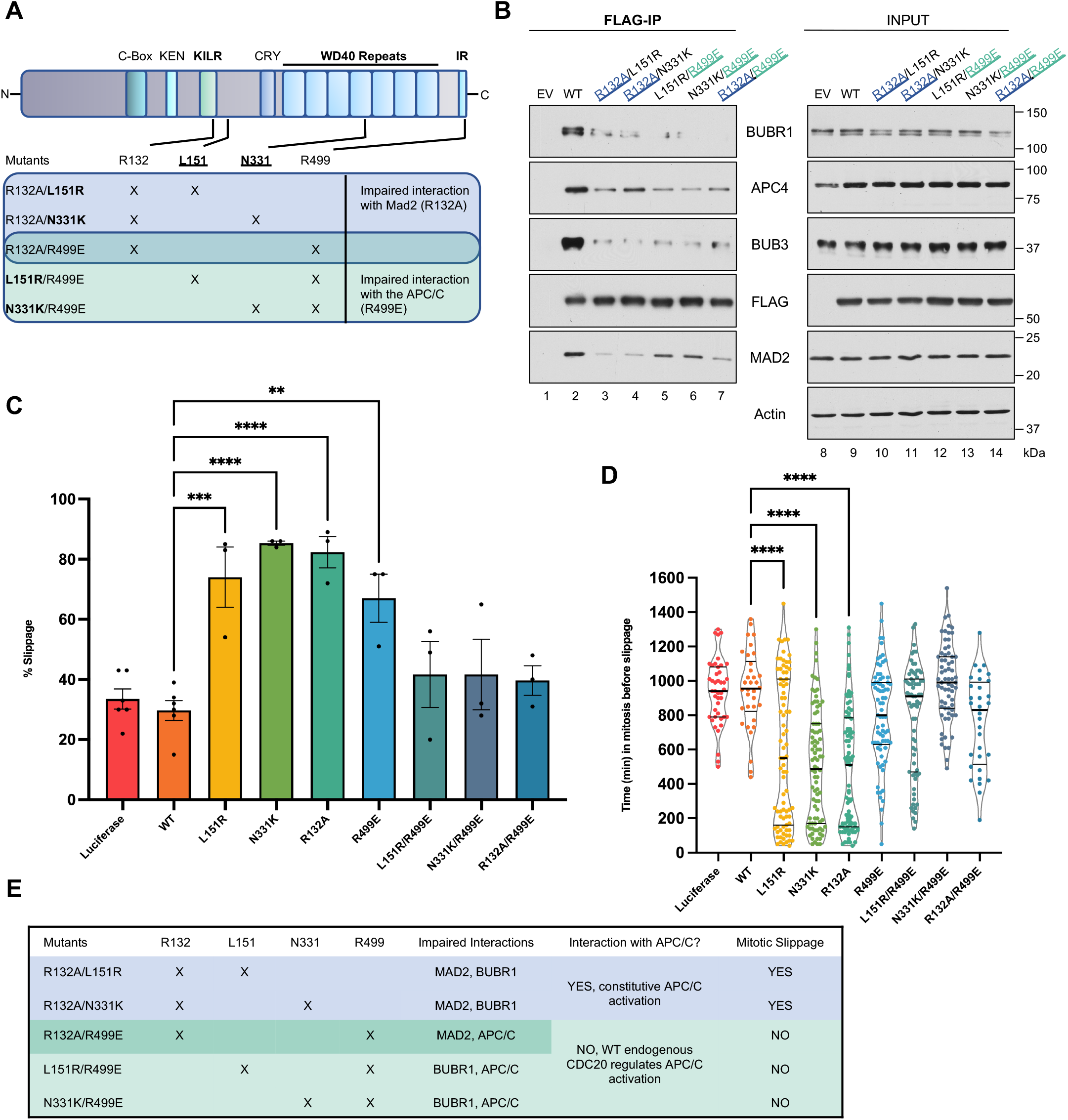
APC/C co-activator activity of familial *CDC20* variants is still required to promote mitotic slippage. **(A)** Schematic showing the different CDC20 protein domains and the locations of the mutations in the different double CDC20 mutant expression vectors tested. The double mutants are classified whether they are MAD2 binding disruptors (with the R132A mutant), APC/C interaction disruptors (with the R499E mutant), or both. **(B)** Immunoblots of different mitotic proteins from FLAG co-immunoprecipitations (co-IPs) on mitotic extracts from cells transfected with FLAG-tagged CDC20 WT or double mutant constructs. Transfected cells were synchronized with thymidine for 22 h, released for 3 h, arrested with nocodazole for 7 h, and harvested. EV: empty vector. **(C)** Quantification of the proportion of cells that underwent mitotic slippage across the different CDC20 mutants from time-lapse experiments performed as in **Figure 3D**. Slippage figures for the single CDC20 mutants (L151R, N331K and N331K) were taken from **Figure 3D** and used here for comparison purposes. An ordinary one-way ANOVA with Dunnett’s multiple comparisons test was used (**: p < 0.01; ***: p < 0.001; ****: p < 0.0001). Error bars: SEM. **(D)** Comparison of the mitotic transit lengths before slippage between the different CDC20 single (from **Figure 3**) and combination mutants. Only cells that underwent mitotic slippage were considered. Cells from a representative experiment were analyzed. Mean slippage times: luciferase, 15 h; WT, 16.2 h; L151R, 9.7 h; N331K, 8.3 h; R132A, 8.1 h; R499E, 13.3 h; L151R/R499E, 13 h; N331K/R499E, 16.5 h; R132A/R499E, 12.7 h. Thick lines indicate the medians of each plot; thin lines indicate the lower and upper quartiles. An ordinary one-way ANOVA with Dunnett’s multiple comparisons test was used (n_luciferase_ = 30; n_WT_ = 35; n_L151R_ = 83; n_N331K_ = 86; n_R132A_ = 89; n_R499E_ = 75; n_L151R/R499E_ = 71; n_N331K/R499E_ = 67; n_R132A/R499E_ = 32; ****: p < 0.0001). **(E)** Summary of the interactions and mitotic activities of the CDC20 double mutants.

### The CDC20 N331K variant exhibits mitotic defects in primary cells from patients and accelerates tumor formation *in vivo*

Although the functional assays performed above suggest that the L151R and N331K mutations exhibit a compromised SAC, we wished to confirm the potential oncogenicity of these mutants in primary patient cells and *in vivo*. We were able to obtain primary skin fibroblasts from the two siblings in fmOGCT7 who are heterozygous for the N331K mutation (**Figure 1B**) and analyzed them by time-lapse microscopy as shown schematically in **Figure 6A**. Interestingly, fibroblasts from both siblings slipped faster through mitotic arrest in the presence of NOC (average times: patient III-1, 3.5 h; patient III-2, 4.2 h) than two control fibroblast lines (average times: Control 1, 8.9 h; Control 2, 11.1 h) (**Figure 6B and C**).

**Figure 6:**
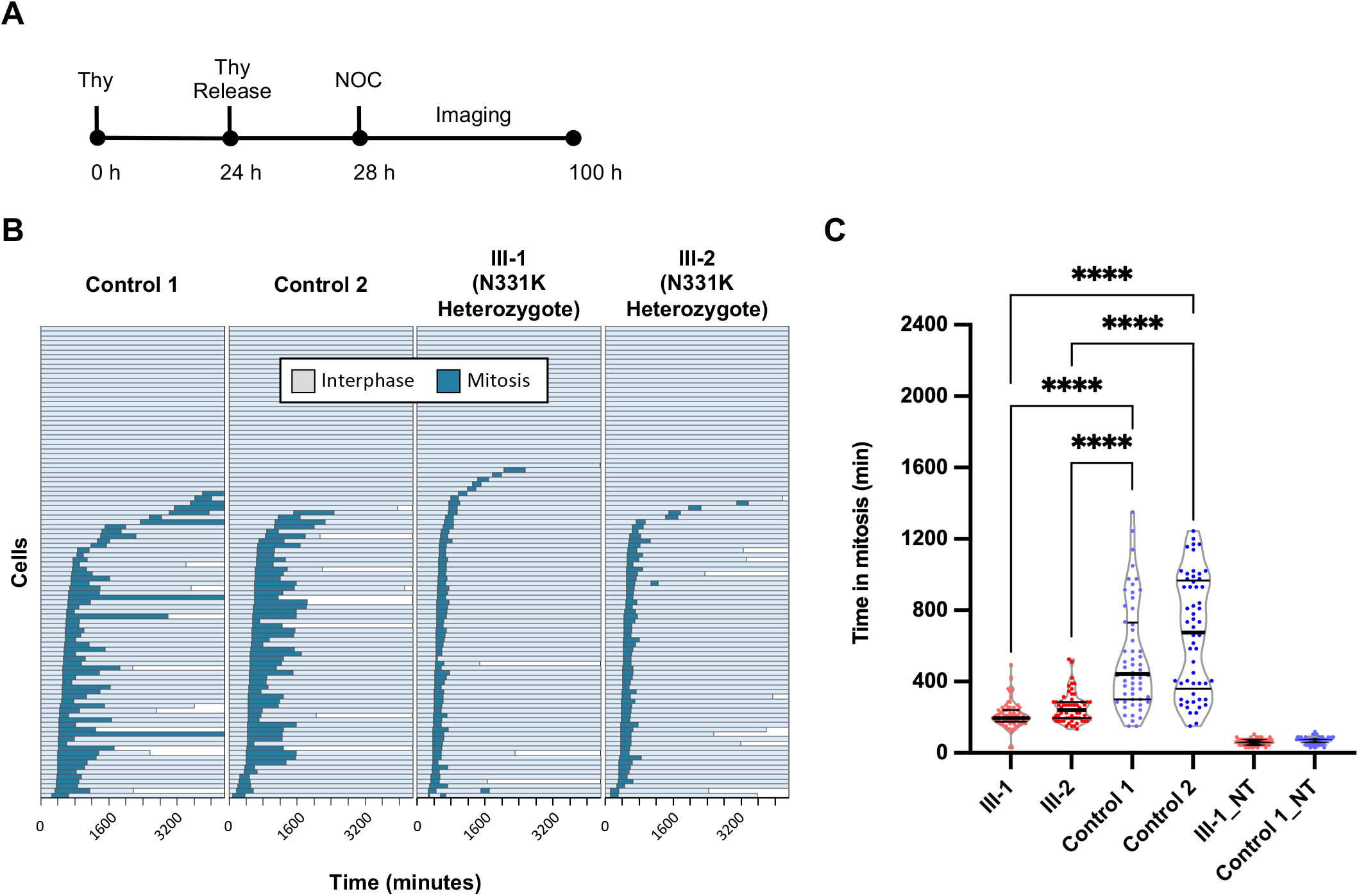
Functional analyses of the CDC20 N331K mutant in patient skin fibroblasts demonstrate a SAC impairment phenotype. **(A)** Visual timeline of the experimental procedure of the mitotic slippage assay on N331K patient skin fibroblasts. **(B)** Visual representation of N331K mitotic skin fibroblasts observed using time-lapse microscopy for 72 h. Fibroblasts were obtained from the two sisters in fmOGCT7 (who are heterozygotes for the N331K variant – III-1 and III-2, see **Fig. 1B**) and WT non-carriers. Images were taken every 15 min following nocodazole treatment. Each analyzed cell was noted for the frame at which it entered mitosis and when it slipped into interphase again. Bar cut-offs into white spaces indicate cell death. The colour scheme is as follows: grey, interphase; blue, mitosis. The frames were converted to and shown in minutes (x-axis). N = 100 cells per patient (y-axis). **(C)** Comparison of mitotic length for patient and control fibroblasts. Only cells that completed mitosis and entered interphase again were considered. NT = not treated with nocodazole. An ordinary one-way ANOVA with Tukey’s multiple comparisons test was used (n_III-1_=70; n_III-2_=63; n_Control 1_=56; n_Control 2_=57; n_III-1_NT_=65; n_Control 2_NT_=72; ****: p < 0.0001). The solid lines indicate the medians of each plot; the dotted lines indicate the lower and upper quartiles.

In order to determine if the N331K mutation is able to drive tumor formation *in vivo*, a knock-in mouse model was derived by introducing the N331K mutation into one allele in C57BL/6 mice using CRISPR-Cas9 gene editing (**Figure 7A, Figure S5**). Mice heterozygous for the N331K mutation (*Cdc20*^N331K/+^) did not significantly develop any more spontaneous tumors than did WT mice (*Cdc20*^+/+^), nor was there a difference in the growth and survival rates between heterozygous and WT animals (**Figure S6A and B**). However, no live births were observed of homozygous (*Cdc20*^N331K/N331K^) animals indicating that this genotype is non-viable (**Figure 7B)**. Moreover, homozygous embryos died between days E4.5 and E10.5 after fertilization and were severely delayed in development compared to WT embryos by the blastocyst stage (**Figure S6C and D**). In support of the observation that the N331K mutation is non-viable, introduction of this mutation in haploid, *S. cerevisiae*, also resulted in lethality (**Figure S7**). Although the N331K variant alone did not result in spontaneous tumorigenesis in heterozygous mice, heterozygous MEFs displayed accelerated mitotic transit compared to WT MEFs under the presence of NOC (averages of 112 min vs. 45 min for WT and mutant, respectively) (**Figure 7C, D and E**). Since the N331K mutation conferred an aberrant mitotic phenotype to MEFs under a pharmacological stress (NOC), we therefore determined if this mutation could enhance tumorigenicity of a system already in an oncogenic context. We crossed our *Cdc20*^N331K/+^ mice with the Eμ-myc cancer model^35,36^ to investigate any cooperation of *Cdc20*^N331K^ with the Eμ-myc onco-transgene, with which mice develop B-cell lymphomas at 2-3 months of age. Eμ-myc^+^ mice carrying one N331K allele (*Cdc20*^N331K/+^/Eμ-myc^+^) developed lymphomas faster and experienced accelerated mortality rates compared to Eμ-myc^+^ mice that were WT for *Cdc20* (*Cdc20*^+/+^/Eμ-myc^+^) (**Figure 7F**). Two separate founder mouse lines carrying the N331K allele were derived and tested in the Eμ-myc model, and no differences in phenotype were found in the experiments mentioned above (data not shown). Our characterization of the N331K variant in patient skin fibroblasts and in mice demonstrate the potential of the germline N331K *Cdc20* mutation to enhance oncogene driven tumors *in vivo*.

**Figure 7:**
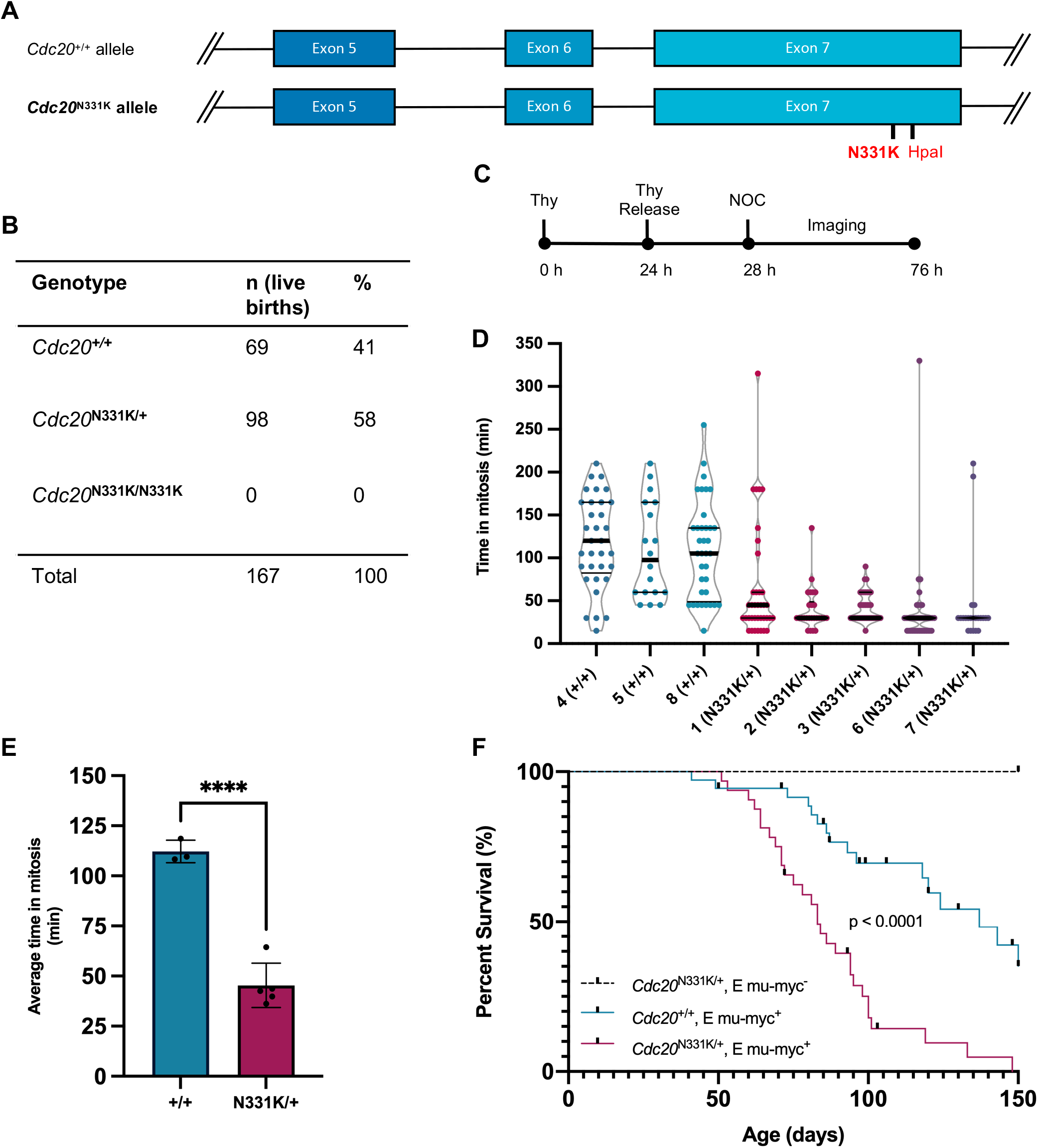
The *Cdc20* N331K mutant exhibit mitotic defects and accelerate tumor formation *in vivo*. **(A)** Schematic of the incorporation of the N331K mutation in the *Cdc20* locus in *Mus musculus* using the CRISPR-Cas9 gene editing technology. An HpaI restriction site was included for genotyping purposes. **(B)** Summary of the genotypes of newborn pups obtained from intercrosses of *Cdc20*^N331K/+^ mice. **(C)** Visual timeline of the experimental procedure of the time-lapse microscopy performed on MEFs extracted from the N331K mouse. **(D)** Scatter violin plots of the duration of mitosis for +/+ (WT) and N331K/+ (het) MEFs from the different isolated mouse embryos. The numbers indicate the MEF IDs. Images were taken every 15 min following nocodazole treatment. Each analyzed cell was noted for the frame at which it entered mitosis and when it entered interphase again. Only cells that completed mitosis and entered interphase again were considered (n_1_ = 37; n_2_ = 26; n_3_ = 36; n_6_ = 41; n_7_ = 22; n_4_ = 33; n_5_ = 18; n_8_ = 36). The thick lines indicate the medians of each plot; the thin lines indicate the lower and upper quartiles. **(E)** Quantification of the average duration of mitosis before slippage between all N331K/+ and WT MEFs analyzed in panel **C**. The individual means in panel **C** were taken and used to compute the average times here. An unpaired, two-tail t test was used (+/+: n = 3, avg. time = 112 min; N331K/+: n = 5, avg. time = 45 min; ****: p < 0.0001). Error bars: SEM. **(F)** Survival (Kaplan-Meier) plot of animals from a cross between N331K heterozygote mice and mice carrying the Eμ-myc onco-transgene. All mice that were observed in moribund states (due to signs of cancer/lymphoma – swollen lymph nodes, hunched posture, roughed fur, hyperventilation, and reduced activity) were euthanized and were considered at endpoint for the calculation of the curves. A log-rank (Mantel-Cox) test was performed to compare the survival curves. *Cdc20*^N331K/+^/Eμ-myc^-^: n=22; *Cdc20*^+/+^/Eμ-myc^+^: n = 36, median survival = 137 d; *Cdc20*^N331K/+^/Eμ-myc^+^: n = 32, median survival = 83 d; p < 0.0001.

## DISCUSSION

Here, we report the identification of two germline *CDC20* missense variants (L151R and N331K) that co-segregate with cancer occurrence in two families. To our knowledge, no germline pathogenic variants have previously been identified in *CDC20* in human cancer. We have shown that the two variants compromise the mitotic checkpoint in various models, accelerating mitotic transit and causing slippage, likely contributing to the tumorigenic process of the various cancers in the families.

The impaired interaction between the familial CDC20 mutants and BUBR1 attenuates the SAC, resulting in more mitotic slippage of cells (**Figure 8**). The reduced affinity of the N331K mutant to BUBR1 is evidenced by the mutation being in the WD40 domain of CDC20, a crucial site of interaction with mitotic substrates that BUBR1 blocks when the SAC is engaged.^37,38^ Although the L151R mutation could not be modelled computationally, we observed that it also had reduced binding to BUBR1, despite not being located in the WD40 domain. This observation could be due to a drastic change from a medium-sized hydrophobic residue (leucine) to one that is large and positively charged (arginine), which could have an impact on protein folding and BUBR1 binding as well. Moreover, a variant similar to the L151R – an L151F mutation – had been identified in a kidney cancer (http://cancer.sanger.ac.uk/cosmic). As had been previously noted,^39^ likely deleterious missense mutations in genes encoding several APC/C proteins have been identified in large-scale sequencing projects and could be selected for, as they result in a “just right” level of genomic instability. At least two *CDC20* missense mutations (L151F and P477A, Table S5 in ref^39^) could fall into this group and are classified as probably damaging by Polyphen-2. Both the L151R and N331K mutants had normal binding to MAD2 during the SAC but their abilities to induce mitotic slippage were comparable to that of the R132A mutant, in which CDC20 is mutated in the MAD2 binding domain (KILR motif). This suggests that impaired CDC20 interactions with BUBR1 and MAD2 have comparable phenotypic outcomes relating to mitotic progression and slippage.

**Figure 8:**
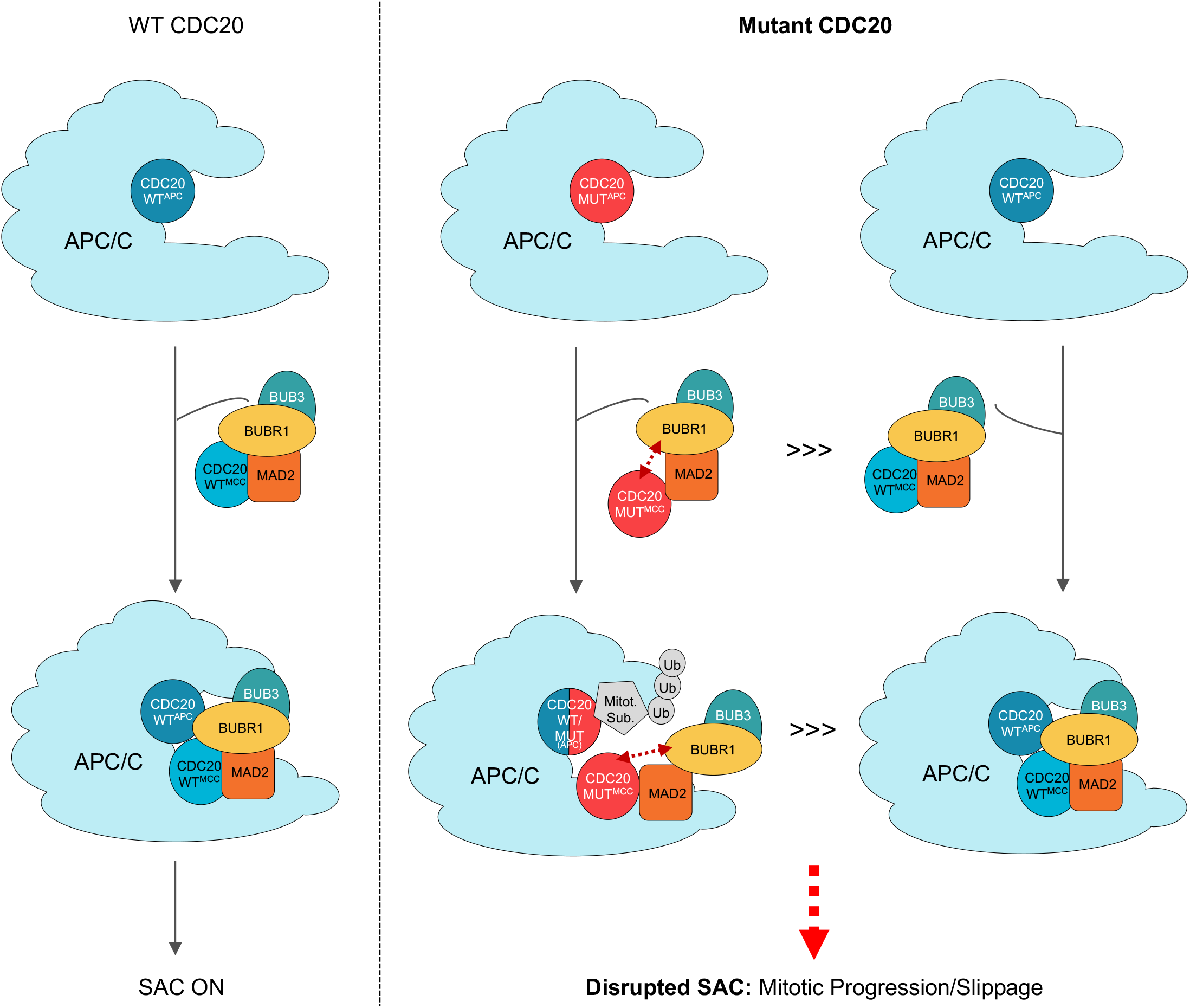
A proposed mechanism by which CDC20 mutants L151R and N331K attenuate the SAC, resulting in aberrant mitotic progression. **Left panel:** During the SAC in cells with WT CDC20, proteins MAD2, BUBR1, BUB3 and CDC20^MCC^ form the mitotic checkpoint complex (MCC) to inhibit the APC/C^CDC20^ complex. The CDC20 molecules that are part of the APC/C or MCC are indicated by their superscripts. **Right panel:** In cells with mutant (L151R or N331K) CDC20, mutant CDC20^MCC^ molecules have an impaired interaction with BUBR1 (shown by red, dotted double arrows). Thus, defective MCCs outcompete the fully functional MCCs (as indicated by >>>), and fewer APC/C^CDC20^ complexes are inhibited during the SAC. Cells experience a disrupted SAC and undergo mitotic slippage

The lack of live births of N331K homozygous mice (*Cdc20*^N331K/N331K^) as well as developmental delays in homozygous embryos corroborate our observation of no homozygous persons in our mOGCT families. The significance of CDC20 in cell and organism viability was likewise reported in the study by Li *et al*.^17^. In their study, mice homozygous for the *Cdc20* allele in which three amino acid residues critical for Mad2 interaction were mutated to alanine (*Cdc20*^AAA^: K129A-R132A-P137A) died at late gestation and embryos of which also exhibited severe developmental delays. Heterozygotes were viable but developed spontaneous tumors at highly accelerated rates, indicating that the SAC-mediated inhibition of Cdc20 is an important tumor-suppressing mechanism. As previously described, the strength of the SAC is crucial in maintaining an orderly mitosis.^12^ A weakened SAC leads to premature chromosomal separation and mitotic slippage.^12^ Conversely, a hyperactive SAC results in lagging chromosomes (leading to monosomies and trisomies) or cell death.^12^ SAC component molecules must also be maintained at specific stoichiometric ratios for proper SAC function. Overexpression of Mad2 in mice leads to aneuploidy and tumorigenesis,^40^ and while mice with reduced levels of Cdc20 do not develop cancer, they accumulate significant amounts of aneuploidy.^41^ At the extreme end of the spectrum, cells without a SAC undergo death in mitosis.^12^ The fact that our heterozygous *Cdc20*^N331K^ mice were viable yet heterozygote MEFs displayed an attenuated SAC under NOC could explain why mutant *Cdc20*^N331K^ molecules alone cannot sustain live homozygous animals – without a WT Cdc20 copy, the SAC is severely weakened, failing to regulate mitosis and maintain normal cell physiology. This suggests that missense mutations in other putative sites of CDC20 are presumably deleterious in development or pathogenic due to the fine balance of CDC20 protein function and appropriate stoichiometric levels necessary for a normal SAC. Indeed, the handful of *CDC20* likely pathogenic or pathogenic variants identified thus far in clinical association studies are potentially linked to problems with germ cell maturation in human infertility.^19–22^ The notion that CDC20 plays a vital role in development could explain the rarity of *CDC20* germline alterations in cancer and why those restricted to tumors are likely to be passenger mutations.

Germline variants in other relevant genes encoding SAC proteins have been associated with an increased cancer risk.^16^ Persons biallelic for pathogenic variants in *BUB1B* (encoding the BUBR1 protein) are at high risk of childhood embryonal tumors such as embryonal rhabdomyosarcoma,^42^ and also gastrointestinal cancers in adults^43^ – both cancers were observed in fmOGCT4. As seen in our *CDC20* variants, pathogenic variants in *BUB1B* lead to SAC impairment. Defects or disruption in the SAC are responsible for an increased rate of aneuploidy during tumorigenesis, but whether this represents a cause or effect of oncogenesis is controversial.^11^ We found subclones with trisomy X and a t(3;13) in blood and skin cells of patients from fmOGCT7, however, such changes can also be seen in individuals without *CDC20* mutations. The significance of these findings is uncertain, but clearly the N331K *CDC20* variant, unlike biallelic *BUB1B* pathogenic variants,^42,43^ does not cause mosaic variegated aneuploidy. Recent studies have shown that low levels of aneuploidy and transient chromosome instability are sufficient to accelerate cancer progression in *p53*-deficient and *Myc*-driven cancer contexts.^44,45^ When we crossed our *Cdc20*^N331K^ mice with the Eμ-myc cancer model, Eμ-myc mice carrying the *Cdc20*^N331K^ allele developed lymphomas and reached clinical endpoint sooner than animals carrying the WT allele of *Cdc20*. The ability of N331K to accelerate *Myc-*driven lymphomagenesis strongly suggests a role of these *CDC20* variants in cancer predisposition. Indeed, there are ongoing efforts to uncover other candidate driver genes in mOGCTs.^46^ We found a D820G mutation in the *KIT* gene in the DG arising in III-2 of fmOGCT7, which was previously identified in hematopoietic neoplasia and testicular germ cell tumors.^47– 49^ A nearby mutation, D816H, was found to play a transforming role in some seminomas via the KIT signal transduction pathway.^50^ Moreover, another study suggested that D820G contributes to drug resistance.^51^ Therefore, while our data argue that the reported *CDC20* variants are predisposing alleles, we cannot conclude that they are full oncogenic drivers as they likely rely on other activating events to initiate and enhance neoplastic transformation.

In summary, we describe cancer-promoting germline variants in *CDC20* in two families ascertained because of the occurrence of familial mOGCTs. These variants dysregulate the mitotic checkpoint. The resulting cancers may not be restricted to the ovaries as pathogenic variants in *CDC20* likely also increase the risk of neoplasia at other sites, as illustrated by the families in this study. Further studies are required to determine the range and potential effects of germline *CDC20* variants and to better understand the mechanisms triggering tumorigenesis through aneuploidy.

## ACKNOWLEDGEMENTS

We thank Drs. Amy Stettner, Blaise Clarke, Ethel Queralt and Rabea Wagener for their help with this project, Mitra Cowan and all personnel at the McGill Integrated Core for Animal Modeling (MICAM) for the derivation of the *Cdc20* N331K mutant mice and the additional experimental support, Dr. Jerry Pelletier for supplying us with the Eμ-myc mice, and Dr. Marcos Malumbres for sharing the conditional *Cdc20* KO MEFs. O.J.C. is a recipient of the Canada Graduate Scholarship – Master’s (CGS-M), the McGill Integrated Cancer Research Training Program (MICRTP) Scholarship, and is supported by the Hilton J. McKeown and George Corcoran Scholarships from McGill University. This work is funded by a Foundation Grant (FDN-48390) of W.D.F. from the Canadian Institutes of Health Research (CIHR), a Project Grant (PJT-156078) of J.G.T. from the CIHR, and a Catalyst Grant (Project No. 160819-001-001) of J.G.T. from the Rare Diseases: Models & Mechanisms Network (RDMM).

## AUTHOR CONTRIBUTIONS

Conceptualization, O.J.C., E.C., W.D.F., and J.G.T.; Investigation, O.J.C., E.C., M.M.-K., J.N., B.R., S.F., Y.W., C.P., L.J., J.C-Z., L.W., A.B., S.B., C.J.R.S., Z.Z., and W.C.C.; Resources, I.G., S.S., D.S., M.T., W.D.F., and J.G.T.; Writing – Original Draft, O.J.C. and E.C.; Writing – Review & Editing, O.J.C., R.S., C.J.R.S., W.D.F. and J.G.T.; Supervision, R.S., C.M.T.G., D.B., J.M., W.D.F., and J.G.T.; Funding Acquisition, W.D.F. and J.G.T.

## DECLARATION OF INTERESTS

The authors declare no competing interests.

## METHODS

### Patients

Germline DNA from blood or saliva was obtained from all available affected members from 5 mOGCT families and sequenced by WES. Additional germline DNA from non-affected family members in fmOGCT4 and fmOGCT7 was obtained to validate *CDC20* mutations and assess co-segregation with the disease. Additional biological samples were collected when possible. Tumoral formalin-fixed paraffin-embedded (FFPE) and fresh frozen (FF) specimens from the mOGCTs and skin primary fibroblasts were also collected from the two sisters in fmOGCT7 (III-1 and III-2). In addition, 85 more FF DNA samples were tested from an anonymized cohort of individuals (16 male and 69 female) with sporadic GCTs. Informed consent was obtained from participants and all analyses were approved by the appropriate ethics committee.

### Whole-exome sequencing (WES) analysis

Genomic DNA from blood and FFPE specimens were respectively subjected to the Agilent 50Mb exon capture kit and the Nextera Rapid Exome kit, following the manufacturers’ protocols. Each sample was sequenced on Illumina HiSeq 2000 (100bp paired-end reads) at the McGill University and Génome Québec Innovation Centre (MUGIC). Reads were first trimmed based on base quality (>30) and were aligned to the human genome (UCSC hg19, NCBI build 37) using BWA (v. 0.5.9).^52^ Then, the Genome Analysis Toolkit (GATK)^53^ performed local realignment of reads around small insertions and deletions (indels) and Picard was used to remove duplicate reads from the final alignment for each sample. GATK was used to ensure an average coverage of 80X of the consensus coding sequence (CCDS) in all samples. Germline variants including single nucleotide variants (SNVs) and indels were called using SAMtools (version 1.8)^54^. Somatic SNVs and indels were called using MuTect^55^ and Indelocator (https://confluence.broadinstitute.org/display/CGATools/Indelocator), respectively, and were then annotated with ANNOVAR. To remove common variants and false positive calls, candidate variants were subjected to several filtering steps and eliminated if they fulfilled any of the following criteria: (i) genomic position of variant covered by <5X and less than 4 reads supporting the alternative variant, (ii) variant has allelic ratio <10% for SNVs or <15% for indels, (iv) variant has allele frequency >0.001 in our non-cancer (>2000 exomes sequenced previously in our centre) or ExAC databases, or variant seen as homozygote in ExAC database. Finally, only the most likely damaging variants (missense, nonsense, splicing [within 6-bp of a splicing junction] and coding indels) that were from highly conserved regions across species and predicted to be disease causing by at least 3 out of 5 bioinformatic software (SIFT, PolyPhen, MutationTaster, Revel and MCAP) were considered for further analysis. The Integrative Genomics Viewer^56^ was used for the manual examination and visualization of all potential candidate variants. We prioritized heterozygous variants from highly conserved regions across species shared among affected family members.

### Simulation study

To estimate the probability of finding a pathogenic mutation/variant in the same gene in exome-sequenced individuals with cancer in 2 out of 5 families, a simulation study was performed. We used the pedigree structures of the families studied and varied two parameters: the number of genes of interest, G, where the presence of a pathogenic variant is assessed, and the frequency of such variants in the general population. The steps taken are explained in detail in **Tables S2A and S2B**.

### Sanger sequencing

All the mutations found by WES were validated by Sanger sequencing using standard protocols. *CDC20* screening on the 85 sporadic GCTs was also performed by Sanger on exons 3 to 6 and 8 to 10, which include important functional motifs. All primers are available upon request.

### Molecular modeling

Coordinates for human CDC20 in complex with the KEN Box of BUBR1 (PDB code 4GGD) were visualized using PyMol.^57^ *In silico* mutation and selection of viable rotamer for the CDC20 K331 residue that prevents any steric clashes was also performed using PyMol.

### Plasmids

*CDC20* cDNA was cloned into the multiple cloning site of the expression vector p3XFLAG-*myc*-CMV-26 (Sigma). Site-directed mutagenesis was used to generate the *CDC20* mutants of interest: L151R was generated through the c.452T>G nucleotide change; N331K through c.993C>A (although fmOGCT7 harbours a c.993C>G change, they both give rise to the same amino acid change and equivalent effects on the mitotic checkpoint have been confirmed in both mutants – data not shown); R132A through c.394_395delinsGC; and R499E through c.1495_1497delinsGAG. The p3XFLAG-*myc*-CMV-26 constructs were used in CDC20 expression and immunoprecipitation experiments. To follow transfected cells through in the time-lapse microscopy experiments, the GFP-expressing bicistronic reporter construct pIRES2-EGFP (BD Biosciences Clontech) was used. The pIRES2-EGFP vector contains an internal ribosomal entry site (IRES), which allows the expression of both CDC20 and GFP proteins independently. WT and *CDC20* mutants were PCR-amplified from the p3XFLAG-*myc*-CMV-26 vector and cloned into the multiple cloning site of pIRES2-EGFP. Double *CDC20* mutants were created with either additional site directed mutagenesis or sub-cloning from the previously made single-mutant constructs. All double mutants were cloned into p3XFLAG-*myc*-CMV-26 and pIRES2-EGFP vectors and tested as described above for the single mutants. All mutations were validated by Sanger sequencing.

### Cell lines, cell culture, transfection, and treatments of eukaryotic cells

HeLa cells and primary fibroblasts were regularly tested for mycoplasma by microscopy inspection following DAPI staining. HeLa cells, primary fibroblasts, and mouse embryonic fibroblasts (MEFs) were cultured using DMEM (Wisent Bioproducts: 4.5 g/L glucose, with L-glutamine, sodium pyruvate and phenol red) with 10% fetal bovine serum (Sigma-Aldrich, Canadian origin) and gentamycin at 50 µg/mL (Wisent Bioproducts). HeLa cells were seeded at 50-60% confluency and Lipofectamine® 2000 (ThermoFisher Scientific) was used for transfection following manufacturer’s instructions, using a 1:2.5 or 1:2 ratio of DNA: Lipofectamine. Cells were treated with the appropriate drugs 24 h post-transfection. To assess mitotic checkpoint function, transfected cells, fibroblasts and MEFs were synchronized with 0.61 mg/mL thymidine (Sigma) for 22-24 h, released for 3-4 h, and treated with 100 ng/mL nocodazole (Sigma-Aldrich) overnight. For the co-immunoprecipitations, mitotic shake-off by dish tapping or strong pipetting of transfected HeLa cells was performed after a 20 h treatment with 9 µM RO3306 (Alexis Biomedicals) released for 75 min in nocodazole, or after thymidine-nocodazole synchronization as described above. *Cdc20* conditional knockout (KO) MEFs (Dr. Marcos Malumbres Lab) were electroporated with the bicistronic *CDC20* constructs using the Gene Pulser Xcell™ Electroporation System (Bio-Rad) according to the manufacturer’s instructions. MEFs were treated with 0.61 mg/mL thymidine and 1 µM 4-hydroxytamoxifen (4-OHT, from Selleckchem) for 22-24 h to synchronize and induce the KO of *Cdc20*. Synchronized cells were released into 4-OHT alone prior to live imaging.

### Western Blotting

Transfected HeLa were harvested and lysed with lysis buffer (50 mM Tris-HCl pH 7.5, 150 mM NaCl, 0.5% NP-40, protease inhibitors tablet from Roche, 50 mM NaF, 10 mM glycerol 2-phosphate disodium salt hydrate, 0.1 mM Na_3_VO_4_ and 1 mM DTT). 40 µg of total protein were separated by SDS-PAGE and probed with antibodies against BUBR1 (BD BioSciences), CDC27 (BD BioSciences), FLAG (Sigma), CDC20 (Santa Cruz), β-actin (Sigma) MAD2 (Covance), and BUB3 (BD BioSciences).

### Cell cycle analysis (Flow cytometry)

Transfected HeLa were harvested and fixed with ice-cold ethanol. Cellular DNA was stained using propidium iodide and analyzed using the FACSCaliburs Analyser (Beckton-Dickinson). Data was analyzed using FlowJo and ModFit softwares.

### Time-lapse microscopy

Cells were synchronized with thymidine for 22-24 h, released for 3-4 h and imaged after the addition of nocodazole. The Zeiss Axiovert 200M microscope was set to take pictures using the brightfield or FITC filter at 10X magnification every 10 or 15 min for transfected HeLa or *Cdc20* conditional KO MEFs with bicistronic CDC20 constructs, mouse embryonic fibroblasts, or patient skin primary fibroblasts. For analysis, 100 cells were followed frame by frame for each cell line. Only green cells were monitored for pIRES2-EGFP-transfected cells. Attached cells were considered to be at interphase and rounded up cells were counted as mitotic. Mitotic slippage times were taken from nuclear envelop breakdown to cell death, slippage, or cytokinesis. Mitotic fates (death or slippage) were determined by cell morphology: cell death was identified by signs of blebbing while slippage was characterized by cells flattening from mitosis without appreciable cytokinesis.

### Immunoprecipitation

Mitotic extracts were quantified and normalized to the same concentration. Approximately 90% of the extract was incubated with anti-FLAG M2 beads (Sigma, EZview RED FLAG M2 Affinity Gel) for 2 h, or with anti-FLAG (Sigma) antibody overnight and then with equilibrated protein G beads (Millipore Sigma, Protein G Agarose, Fast Flow) for 2h. After several washes with lysis buffer, samples were resuspended in 4X loading buffer (200 mM Tris-HCl, 4% SDS, 40% glycerol, 4% β-mercaptoethanol, 0.012% bromophenol blue and 50 mM DTT) and resolved by SDS-PAGE. Mouse TrueBlot ULTRA (anti-Mouse Ig (Rockland) secondary antibody) was used to probe for FLAG and CDC20 primary antibodies.

### OncoScan

Tumor samples from fmOGCT7 (III-1 mixed GCT and III-2 DG) were treated with the OncoScanTM FFPE Assay Kit then subjected to the OncoScanTM array. The Nexus Express Software for OncoScan v.3.1 (BioDiscovery, El Segundo, California) was used to call copy number (CNV) and allelic imbalance (AI) across the genome. Briefly, the Fast Adaptive States Segmentation Technique (FASST2) algorithm (a Hidden Markov Model based approach) segments copy number regions based on median log-ratio derived from all probes in a given region. The segments were detected with a significance threshold of 1.0^-8^, a minimum of 10 probes per segment, and a maximum probe spacing of 1000 kbp between adjacent probes. Then, the log_2_ ratio was considered for the segments’ classifications as follows: gains (> 0.08), losses (< -0.1), high copy gains (> 0.7) and big losses (< -1.1). The AI state was determined for each segment based on the median B-allele frequency (BAF) of heterozygous makers. Similar AI results were obtained with WES data using ExomeAI (data not shown).^58^

### Immunohistochemistry

Immunohistochemistry (IHC) was performed at the Segal Cancer Centre Research Pathology Facility (Jewish General Hospital). Tissue samples were cut at 4 µm, placed on SuperFrost/Plus slides (Fisher) and dried overnight at 37°C, before IHC processing. The slides were then loaded onto the Discovery XT Autostainer (Ventana Medical System). All solutions used for automated IHC were from Ventana Medical Systems unless otherwise specified. Slides underwent de-paraffinization, heat-induced epitope retrieval (CC1 prediluted solution Ref: 950-124, standard protocol). Immunostaining for p53 and CDC20 was performed online using a heat protocol. Briefly, a pre-diluted mouse monoclonal anti-p53 (Clone Bp53-11, Ventana Medical Systems, Inc.) and a rabbit polyclonal anti-CDC20 (Abcam #ab72202) at 1:300 were manually applied for 32 min at 37°C, followed by the appropriate detection kit (OmniMap anti-Mouse-HRP, Ref: 760-4310 and OmniMap anti-Rabbit-HRP, Ref: 760-4311) for 8 min, followed by ChromoMap-DAB (Ref: 760-159). The omission of the primary antibody was included as a negative control. Slides were counterstained with hematoxylin for 4 min, blued with Bluing Reagent for 4 min, removed from the autostainer, washed in warm soapy water, dehydrated through graded alcohols, cleared in xylene, and mounted with Eukitt Mounting Medium (EMS, Ref: 15320). Slides were analyzed by conventional light microscopy (or scanned).

### Fluorescence *in situ* hybridization

Fluorescence *in situ* hybridization (FISH) analyses were performed on paraffin slides with tumor tissue from affected individuals from the mOGCT families as previously described.^59^ For the detection of chromosomal aneuploidies, commercially available chromosome enumeration probes for chromosomes 4, 10 and 20 (Abbott) were applied. In the familial cases, the locus specific LSI RUNX1/RUNX1T1 (8q21.3;21q22) Dual Color Dual Fusion probes (Abbott) mixed with a chromosome enumeration probe for chromosome 18 (Abbott) were used. The FISH results on the tissue microarrays (TMAs) were evaluated using Zeiss fluorescence microscopes equipped with the appropriate filter sets. For digital image acquisition and processing, the ISIS digital image analysis system (MetaSystems, Altlussheim, Germany) was applied.

### Generation of the *Cdc20*^N331K^ mouse model

The GRCm38.p4 assembly from the NBCI database was used to map the *Cdc20* genomic sequence in *Mus musculus*. A single-stranded guide RNA (sgRNA) sequence (ATGATAACATTGTCAACGTG) and a single-stranded deoxy-oligonucleotide (ssODN) donor repair template (GCCTTGATGTTGAGTGAATGTTTGCAGGGGAGCCCATCCACTTTCTCCAGGACCACT AGGCCACACGTTAACAATCTTATCATTGCCACCACTTGCCAGATGTCGTCCATCTGGGGCCCAGCGAAGC) were used to respectively target the specific *Cdc20* locus and introduce the c.993C>G, p.N331K mutation and a silent base substitution for an HpaI site. The sgRNA (Alt-R® CRISPR-Cas9 crRNA and tracrRNA), recombinant Cas9 protein (Alt-R® S.p. Cas9 Nuclease V3) and the ssODN (Ultramer DNA Oligo) were all purchased from Integrated DNA Technologies. Microinjections (50 ng/μl Cas9, 50 ng/μl sgRNA, 30 ng/μl ssODN and 100 mM KCl) were performed on 1 side of fertilized C57BL/6 embryos (Charles River) at 1.5 dpc. Injected embryos were subsequently implanted in pseudo-pregnant females. All microinjections were performed by personnel at the McGill Integrated Core for Animal Modeling (MICAM). Mosaic CRISPR-edited mice were screened with targeted deep sequencing (Illumina MiSeq PE250) using the following primer sequences: ACACTGACGACATGGTTCTACAAGTCCCTCGTTTATAGCTGAG and TACGGTAGCAGAGACTTGGTCTACCATGATGTTCGGGTAGC. MiSeq results were analyzed using the Integrative Genomics Viewer (IGV) software.^56^ Founders were obtained and backcrossed with WT C57BL/6 mice. Genotyping was performed using restriction fragment length polymorphism analysis: PCR was performed (with primers ACAATTAAACTCAACCCCGCC and GAATGGACCAGTCCAACAAAGG) and amplicons were digested with the HpaI restriction enzyme. Genotypes were confirmed by Sanger sequencing. *Cdc20*^N331K^ mosaic founders were generated by personnel in the McGill Integrated Core for Animal Modeling (MICAM). Two separate founder lines were maintained throughout the study (see **Figure S5** for more details). Animals were kept in specific-pathogen-free (SPF) rooms. All animal studies were performed in accordance with ethical guidelines and protocols approved by our research institute.

### Primary mouse embryonic fibroblasts

Pregnant females from *Cdc20*^N331K/+^ heterozygote intercrosses were euthanized 13.5 dpc. Embryos were harvested, minced, digested in 0.25% trypsin/EDTA (Wisent Bioproducts) containing 10 mM HEPES (Wisent Bioproducts) at 37°C, and filtered through a 100-μm cell strainer. MEFs were maintained at 37°C in DMEM (Wisent Bioproducts: 4.5 g/L glucose, with L-glutamine, sodium pyruvate and phenol red) with 10% fetal bovine serum (Sigma-Aldrich, Canadian origin). MEFs were subjected to the time-lapse microscopy experiments as described above.

### Survival and tumor studies in mice

*Cdc20*^N331K/+^ and *Cdc20*^+/+^ mice were monitored once a week for any signs of disease and cancer. Mice were euthanized at clinical endpoint (i.e., presence of tumors, inactivity, etc.). The weights of newborns were taken weekly for the first 9 weeks after birth. To study the viability of *Cdc20*^N331K/N331K^ homozygote embryos, zygotes from heterozygote crosses were harvested from females at 0.5 dpc and cultured in EmbyroMax® Advanced KSOM media (Sigma). For tumor formation studies, *Cdc20*^N331K/+^ mice were crossed with the Eμ-myc cancer model (Dr. Jerry Pelletier Lab). Mice were monitored weekly and euthanized when in moribund states and/or at clinical endpoint (signs of cancer/lymphoma – swollen lymph nodes, hunched posture, inactivity).

### Checkpoint cell survival assay in *S. cerevisiae*

The endogenous copy of *S. cerevisiae* CDC20 was replaced using KanMX in transformed yeast carrying a duplicate copy of CDC20 in a URA3 plasmid. Then, a CDC20 copy with the N407K mutation (equivalent to N331K in humans) or the well-known CDC20 KEN mutant (N405A/N407A/Q473A/R517L)^5^ was transformed using the LEU selection marker. The CDC20 WT URA3 plasmid was driven out by 5FOA treatment. The cell survival assay was performed at several dilutions yeast using 50 μg/ml nocodazole for up to 4 h.

### Statistical analysis

Statistical analysis was performed with the GraphPad Prism (Version 9) software (GraphPad, San Diego, CA). For comparisons of more than 2 groups in the functional assays, an ordinary one-way ANOVA was performed followed by post-hoc pairwise comparisons (Dunnett’s or Tukey’s multiple comparisons test). A p-value < 0.05 was considered statistically significant. For data presented in scatter-violin plots, the median and upper and lower quartiles were considered. For histograms, data were expressed as mean ± standard error of the mean (SEM) or mean ± standard deviation (SD). All other statistical analyses are described in the relevant table or figure legends in the Supplemental Information.

## Notes

### Competing Interest Statement

The authors have declared no competing interest.

