## Supplemental Information for "Germline Missense Variants in *CDC20* Result in Aberrant Mitotic Progression and Familial Cancer"

**Table S2A: Estimated probability of finding two or more families where all sequenced individuals carry a variant in the same gene.** The number of variants included in the simulations can be seen in **Table S2B** (see also **Table S1A**). Estimates are based on 5000 simulations

| Allele frequency threshold | Number of genes of interest, $G$ | | | |
|---|---|---|---|---|
| | 25,494 | 10,000 | 1,000 | 500 |
| 0.001 | **0.596** | **0.307** | **0.032** | **0.01** |
| 0.001 disease causing | **0.201** | **0.079** | **0.005** | **0.001** |
| 1e-4 | **0.146** | **0.056** | **0.002** | **8e-4** |
| 1e-5 | **0.041** | **0.022** | **6e-4** | **<1/5000** |
| 1e-6 | **0.035** | **0.009** | **2e-4** | **<1/5000** |

**Table S2B: Numbers of variants, $M_f$, passing filtering criteria used for the simulations.** Variants had at least 5X coverage, at least 4 alternative alleles, were restricted to be either nonsense mutations, missense mutations, splice site variants, or coding indels. Number of variants passing MAF filters are shown, as well as the number also predicted to be disease causing by at least three of five algorithms.

| **Minor allele frequency** | **fOGCT1** | **fOGCT3** | **fOGCT4** | **fOGCT5** | **fOGCT7** |
|---|---|---|---|---|---|
| **<0.001** | 192 | 165 | 253 | 244 | 147 |
| **<0.001 disease causing** | 104 | 88 | 147 | 116 | 74 |
| **<1e-4** | 81 | 75 | 127 | 93 | 63 |
| **<1e-5** | 45 | 41 | 83 | 44 | 45 |
| **<1e-6** | 39 | 36 | 74 | 38 | 37 |

To estimate the probability of finding a pathogenic mutation/variant in the same gene in exome-sequenced individuals with cancer in two of five families, we performed a simulation study. We used the pedigree structures of the families studied (**Figure 1A and B**, and **Figure S1**), and varied two parameters: the number of genes of interest, $\boldsymbol{G}$**,** where the presence of a pathogenic variant is assessed, and the frequency of such variants in the general population. The steps taken were as follows:

1. The number of genes with coverage at least 5X and with good quality results from the WES analysis was identified for each individual (**Table S1**). For the simulation, we assumed that the number of such good quality genes was the same for all sequenced individuals, and was equal to the maximum value observed: 25,494.
2. We also identified different subsets of genes of interest, of size $\boldsymbol{G}$**.** These subsets could correspond to genes that are likely candidates for cancer. (See Note 1 below).
3. By varying an allele frequency threshold filter, the number of WES variants eligible for consideration varied (**Table S2B** and Note 2 below).

4. We defined $\boldsymbol{M_f}$ as the maximum number of eligible variants found in members of sequenced family $f$, for a given allele frequency threshold.
5. These $\boldsymbol{M_f}$ variants were randomly transmitted through the families following Mendelian rules, assuming independence of transmission for each mutation (see Note 3 below).
6. We then identified the $\boldsymbol{L_f}$ variants, a subset of $\boldsymbol{M_f}$, that were shared among all sequenced individuals within family $f$.
7. These $\boldsymbol{L_f}$ variants were randomly assigned, with replacement, to each of the 25,494 genes with good coverage. This was done separately for each family.
8. Steps 4-6 were repeated 5000 times for each allele frequency threshold and for each value of $\boldsymbol{G}$. We then calculated the proportion of the 5000 simulations where the same gene, within gene subset $\boldsymbol{G}$**,** was randomly assigned one of the $\boldsymbol{L_f}$ variants in two or more families; these results are in **Table S2A**.

Note 1: The number of genes considered, $\boldsymbol{G}$**,** was varied from a maximum equal to the number of genes sequenced with good quality (25,494), to several smaller values that might be relevant for cancer pathways: such as 10,000, 1000, and 500.

Note 2: The variant allele frequency filter was varied from 0.001 to 1e-6. Filtering of variants after WES included a step based on the allele frequencies seen in the EXAC database. However, for mutations that were not seen in this database, it is impossible obtain a good estimate of the allele frequency. The number of mutations or variants used in our simulations varied with the allele frequency threshold as seen in **Table S2B**.

Note 3: Within families, the following rules were applied to randomly generate transmission patterns following Mendelian rules: For each family, we randomly assigned $\boldsymbol{M_f}$ variants to one key individual, then randomly transmitted this allele to other individuals in the family following Mendelian rules.
fmOGCT1. $M_1$ variants are assigned to the mother and dropped to the daughter with probability 0.5.
fmOGCT3. $M_3$ variants are assigned to the mother/aunt. Each variant was dropped to the daughter with probability 0.5 and to the niece with probability 0.25.
fmOGCT4. $M_4$ variants are assigned to the mother. For each of the $M_4$ variants, they are dropped to each of the 3 sequenced children with probability 0.5.
fmOGCT5. $M_5$ variants are assigned to the mother and dropped to each daughter with probability 0.5.
fmOGCT7. $M_7$ variants are assigned to one of the sisters. Using probability 0.25, each variant is randomly assigned to the other sister.

### Table S3: WES analysis on tumors

| fOGCT7_III-1 mixed GCT | | | | | | | | | | | | | | | | |
|---|---|---|---|---|---|---|---|---|---|---|---|---|---|---|---|---|
| Gene | gPosition | VarConsequence | BaseRef | BaseAlt | VarType | altReads | TotalReads | Amino_acids | 1KMAF | EVS_MAF | SIFT | Polyphen2 | MutationTaster | Revel | MCAP | COSMIC_ID | COSMIC_OCCUR |
| AKAP1 | chr17:55191929 | nonsynonymous SNV | G | T | het | 5 | 47 | p.S738I | 0 | 0 | 0.01,0.99,D | 0.984,D | 1,1.0,D | 0,42 | 0,016367164 | | |
| AKT2 | chr19:40742002 | nonsynonymous SNV | C | T | het | 21 | 69 | p.D262N | 0 | 0 | 0.01,0.99,D | 0.975,D | 1,1.0,D | 0,277 | 0,03189704 | | |
| ITSN2 | chr2:24428145 | nonsynonymous SNV | C | A | het | 8 | 69 | p.G1540V | 0 | 0 | 0,1.00,D | 0.656,P | 1,1.0,D | 0,248 | 0,051187172 | | |
| PLXNC1 | chr12:94542926 | nonsynonymous SNV | C | A | het | 4 | 22 | p.A60E | 0 | 0 | 0.01,0.99,D | 0.705,P | 1,1.0,D | 0,233 | 0,846793339 | | |
| PSMB10 | chr16:67970130 | nonsynonymous SNV | G | T | het | 8 | 51 | p.A77D | 0 | 0 | 0,1.00,D | 0.992,D | 1,1.0,D | 0,498 | 0,020379166 | | |
| RDH5 | chr12:56115092 | nonsynonymous SNV | C | T | het | 6 | 56 | p.R42C | 0 | 0 | 0.03,0.97,D | 0.964,D | 0.912,0.912,D | 0,708 | 0,309622428 | | |
| fOGCT7_III-2 dysgerminoma | | | | | | | | | | | | | | | | | |
| Gene | gPosition | VarConsequence | BaseRef | BaseAlt | VarType | altReads | TotalReads | Amino_acids | 1KMAF | EVS_MAF | SIFT | Polyphen2 | MutationTaster | Revel | MCAP | COSMIC_ID | COSMIC_OCCUR |
| EFEMP1 | chr2:56108849 | nonsynonymous SNV | C | G | het | 17 | 109 | p.G180R | 0 | 0 | 0.19,0.81,T | 1.0,D | 0.905,0.905,D | 0,601 | 0,147918891 | | |
| KIT | chr4:55599333 | nonsynonymous SNV | A | G | het | 68 | 114 | p.D820G | 0 | 0 | 0,1.00,D | 1.0,D | 1,1,A | 0,894 | 0,160209446 | COSM1316 | 2(testis),5(haematopoietic_and_lymphoid_tissue) |

Note: Highlighted in yellow are putative driver mutations in each of the tumors analyzed.

**Table S4: Fluorescence *in situ* hybridization on available tumors**

FISH analyses were done on available tumors using 6 probes complementary to different chromosomes. Results are shown as number of copies found for each probe and the proportion of cells showing the fluorescence. CEP, chromosome enumeration probes; MGCT, mixed GCT.

| | | CEP4 | CEP10 | CEP20 | 8q21.3RUNX1T1 | 21q22RUNX1 | CEP18 |
|---|---|---|---|---|---|---|---|
| **fmOGCT7** | **Dysgerminoma (III-2)** | 3-8 (78% nuclei) | ne | ne | 3-6 (70% nuclei) | 3-7 (70% nuclei) | 3 (14% nuclei) |
| | **MGCT (III-1)** | 3-5 (17% nuclei) | ne | ne | 3-5 (30% nuclei) | 3-10 (30% nuclei) | 3-5 (30% nuclei) |

**Table S5: CDC20 and P53 IHC staining on available tumors**

For CDC20 IHC staining the proportion of positive cells and the intensity of the staining was evaluated using a semi-quantitative method. Proportion values correspond to ranges of percentage of stained cells: 1 (1-10%), 2 (11-25%), 3 (26-50%), 4 (51-75%) and 5 (76-100%). Intensity values of CDC20 staining are as follows: 0 (none), 1 (weak), 2 (moderate) and 3 (strong). GB, gonadoblastoma (usually in association with dysgerminoma); YST, yolk sac tumor; IT, immature teratoma; and MGCT, mixed germ cell tumor.

| | **CDC20** | | **P53** |
|---|---|---|---|
| | **Proportion** | **Intensity** | |
| GB (n=6) | 4,8 (SD=0,4) | 3 (SD=0) | |
| Dysgerminoma (n=13) | 4,5 (SD=1,4) | 2,2 (SD=1) | |
| YST (n=7) | 3,9 (SD=0,8) | 2,1 (SD=0,5) | |
| IT (n=11) | 3,7 (SD=0,8) | 1,8 (SD=0,6) | |
| MGCT (n=9) | 3,1 (SD=1,3) | 1,5 (SD=0,8) | |
| MGCT (III-1, fmOGCT7)-YST | 4 | 2 | WT |
| MGCT (III-1, fmOGCT7)-Mature | 3 | 2 | WT |
| Dysgerminoma (III-2, fmOGCT7) | 4 | 3 | WT |
| Melanoma (I-2, fmOGCT4) | 5 | 2 | WT |

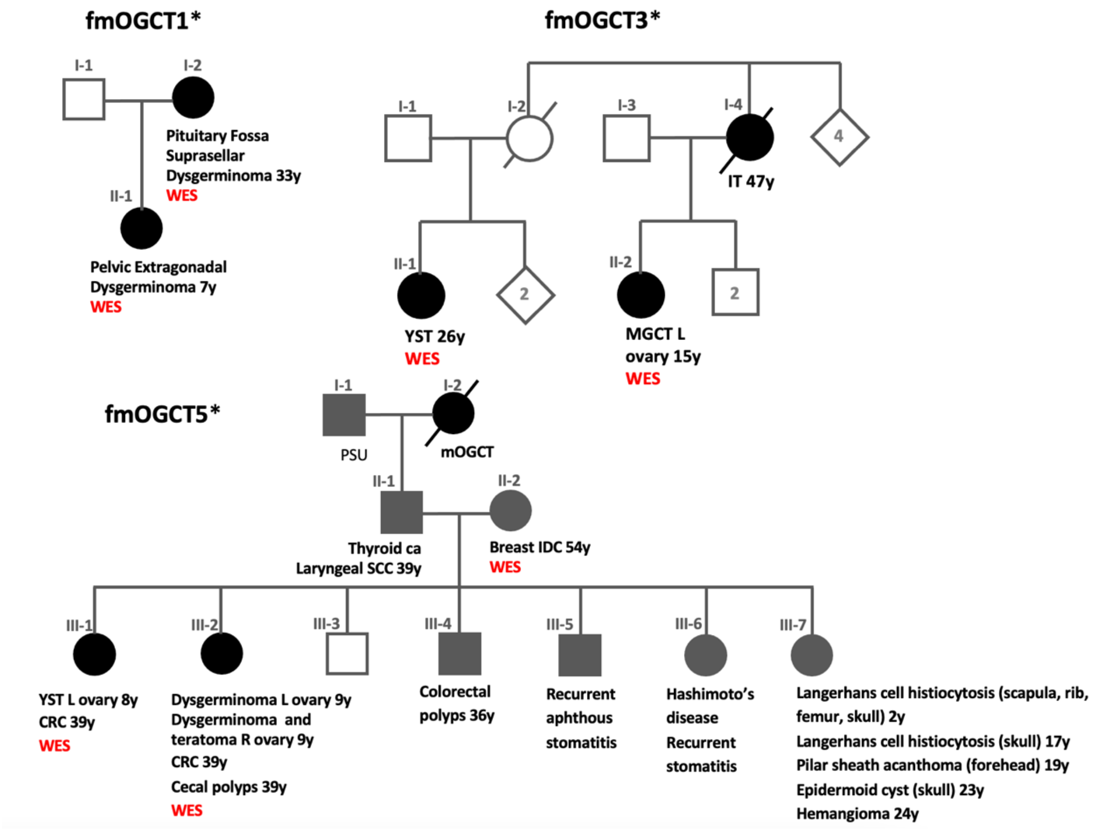


**Figure S1: mOGCT families with no CDC20 mutations found.**
Here we present the revised pedigrees from three families with aggregation of mOGCT (black-filled circles) and other malignancies in some cases (grey-filled shapes) that underwent whole-exome sequencing. Sequenced individuals are indicated by 'WES' under their diagnosis. No *CDC20* mutation was found in any of them. IT, immature teratoma; YST, yolk sac tumor; MGCT, mixed mOGCT; PSU, primary site undetermined; SCC, squamous cell carcinoma; IDC, invasive ductal carcinoma; L, left; R, right; CRC, colorectal cancer.

*These families were first described by the following authors:

fmOGCT1: Chisholm JC; Darmady JM; Kohler JA. Dysgerminoma in Mother and Daughter: Use of Lactate Dehydrogenase as a Tumor Marker in the Child. Pediatric Hematology and Oncology. 1995

fmOGCT3: Stettner AR, Hartenbach EM, Schink JC, Huddart R, Becker J, Pauli R, et al. Familial Ovarian Germ Cell Cancer: Report and Review. American Journal of Medical Genetics. 1999

fmOGCT5: Mandel M, Toren A, Kende G, Neuman Y, Kenet G, Rechavi G. Familial Clustering of Malignant Germ Cell Tumors and Langerhans' Histiocytosis. Cancer. 1994

**A**

| | fOGCT1 | | fOGCT3 | | fOGCT4 | | | | fOGCT5 | | | fOGCT7 | |
|---|---|---|---|---|---|---|---|---|---|---|---|---|---|
| **Sample name** | I-2 | II-1 | II-2 | II-1 | I-2 | II-2 | II-6 | II-4 | II-2 | III-2 | III-1 | III-1 | III-2 |
| # all called variants | 227K | 256K | 234K | 226K | 164K | 165K | 160K | 166K | 218K | 219K | 201K | 167K | 85K |
| #shared variants with 5X coverage and at least 4 alt | 110964 | | 134056 | | 69978 | | | | | 21331 | | 50087 | |
| # missense, nonsense, splicing, coding indels | 8282 | | 6838 | | 6299 | | | | | 1538 | | 6574 | |
| # MAF < 0.001 in EXAC and in-house Exomedb and not seen as homozygous in EXAC | 77 | | 22 | | 41 | | | | | 76 | | 49 | |
| # predicted to be disease causing in 3 out of 6 software and manual inspection | 44 | | 10 | | 23 | | | | | 29 | | 21 | |

**B**

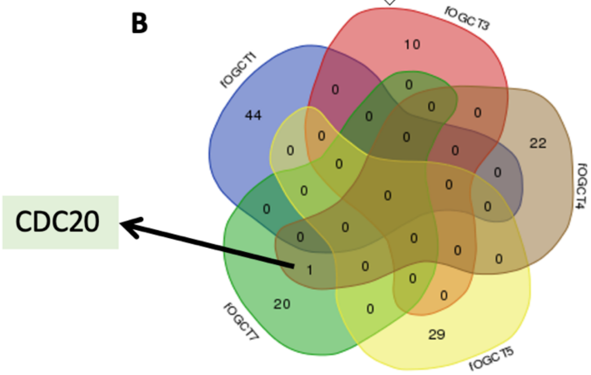


**Figure S2: Whole exome sequencing pipeline chart analysis in familial mOGCT.**
**(A)** Five of the 8 reported families with mOGCT inheritance were studied by WES. After several filtering analyses, rare variants shared by at least two families were assessed.
**(B)** Only the *CDC20* gene harboured putative pathogenic variants in two families.

| | | fmOGCT4 | | | | fmOGCT7 | | | |
|---|---|---|---|---|---|---|---|---|---|
| Genomic position | Mutation | I-2 | II-2 | II-4 | II-6 | III-1 | III-1 mixed GCT | III-2 | III-2 dysgerminoma |
| chr1:43825664 | c.452T>G [p.L151R] | het (138/269) | het(116/227) | het(96/207) | het(100/186) | - | - | - | - |
| chr1:43826548 | c.993C>G [p.N331K] | - | - | - | - | het(26/65) | het(40/86) | het(4/7) | het(28/57) |

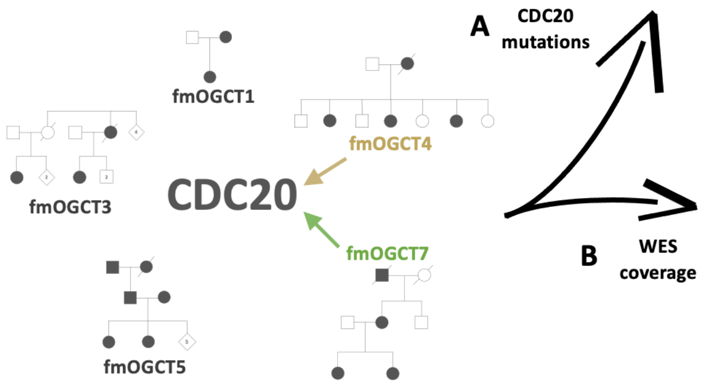


| Family | Individual | Mean coverage | %CCDS bases >=5x coverage | %CCDS bases >=10x coverage | %CCDS bases >=20x coverage | %CCDS bases >=50x coverage |
|---|---|---|---|---|---|---|
| fmOGCT4 | I-2 | 127.24 | 96 | 93.5 | 89.4 | 78.2 |
| fmOGCT4 | II-2 | 129.53 | 95.8 | 93.2 | 89 | 77.6 |
| fmOGCT4 | II-4 | 130.06 | 96 | 93.4 | 89.4 | 78.3 |
| fmOGCT4 | II-6 | 117.66 | 95.9 | 93.2 | 88.9 | 76.1 |
| fmOGCT7 | III-1 | 132.7 | 96 | 93.5 | 89.5 | 78.5 |
| fmOGCT7 | III-2 | 17.75 | 87.1 | 71.9 | 37.7 | 4.1 |
| fmOGCT7 | III-1 mixed GCT | 66.98 | 97.8 | 96.4 | 91 | 58.2 |
| fmOGCT7 | III-2 DG | 47 | 97 | 94 | 81.2 | 36.6 |

**Figure S3: Whole exome sequencing data in *CDC20* mutation-carrier families.**

**(A)** Variant c.435T>G [p.L151R] was shared by four affected women in fmOGCT4 and conserved in a colorectal cancer (CRC) from one of them (data not shown). Variant c.993C>G [p.N331K] was shared by two affected siblings of fmOGCT7, and again conserved in their mOGCT tumors. In parenthesis we found number of altered vs total reads in these positions from the WES analyses.

**(B)** The coverage obtained from WES in the studies families. Only two samples showed low coverage (fmOGCT7, III-2 and fmOGCT7, III-2 DG). DG dysgerminoma, CCDS consensus coding DNA sequence.

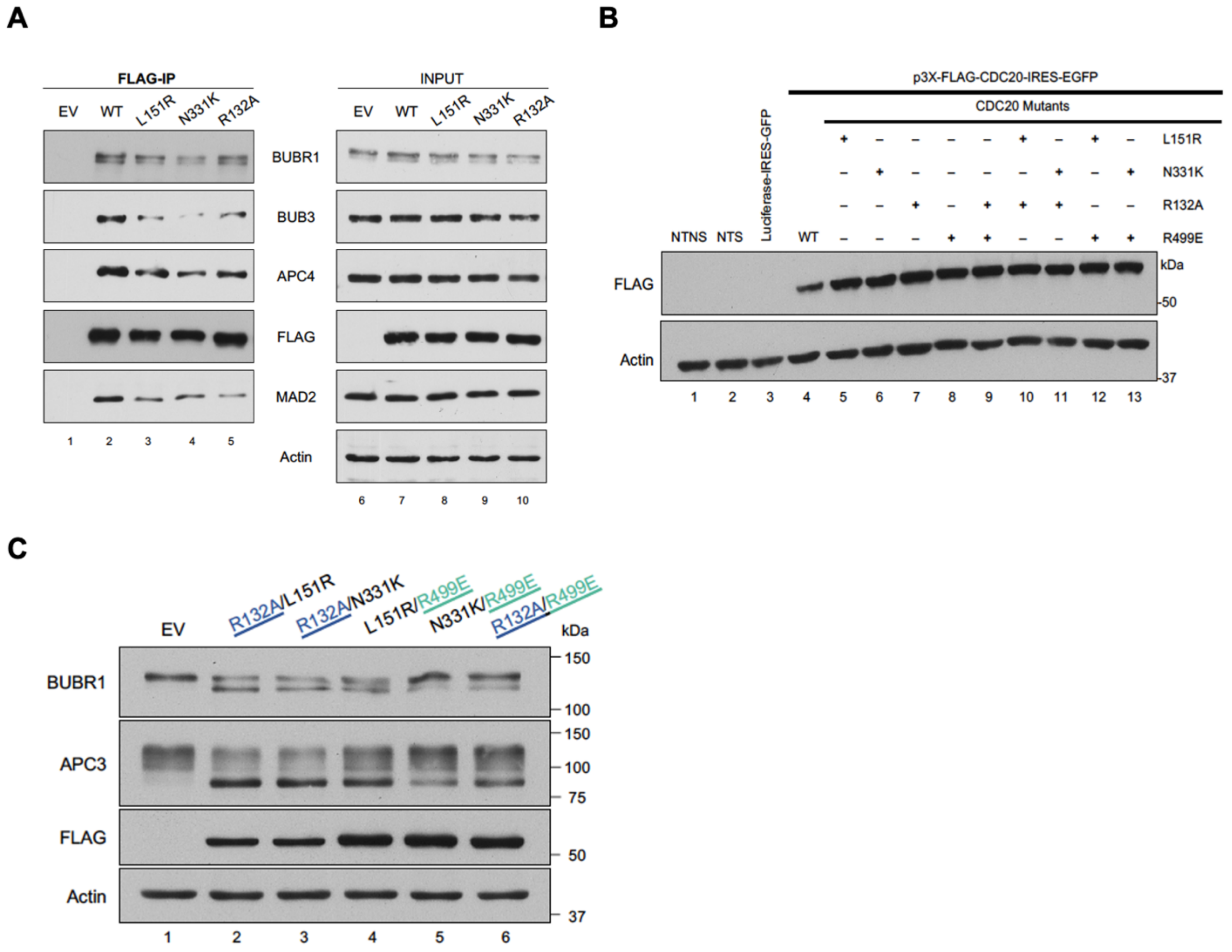


**Figure S4: Immunoblots of mitotic extracts transfected with different CDC20 mutant constructs**

**(A)** Co-immunoprecipitation (Co-IP) reactions of FLAG-tagged CDC20 mutants from thymidine-nocodazole synchronized HeLa cells show reduced CDC20-BUBR1 affinity. FLAG-tagged WT or mutant CDC20-expressing constructs were transfected into HeLa cells, synchronized with thymidine, arrested in mitosis with nocodazole, and harvested. Co-IPs were performed on the extracts, and proteins of which were resolved using SDS-PAGE and probed with the appropriate antibodies. Compared to the WT, the CDC20 variants from the mitotic extracts have reduced affinity to BUBR1, BUB3 and MAD2.

**(B)** Expression blots of HeLa cells transfected with the various CDC20 mutants cloned into the bicistronic pIRES-EGFP expression plasmid. Transfected cells were synchronized with thymidine and nocodazole, harvested as in (A). NTNS: non-transfected, non-synchronized; NTS: non-transfected, synchronized.

**(C)** Expression blots of extracts of cells transfected with FLAG-tagged CDC20 double mutant constructs, synchronized with thymidine for 22 h, released for 3 h, and arrested with nocodazole for 16 h. The phosphorylated forms of BUBR1 and APC3 are used as mitotic markers. EV: empty vector.

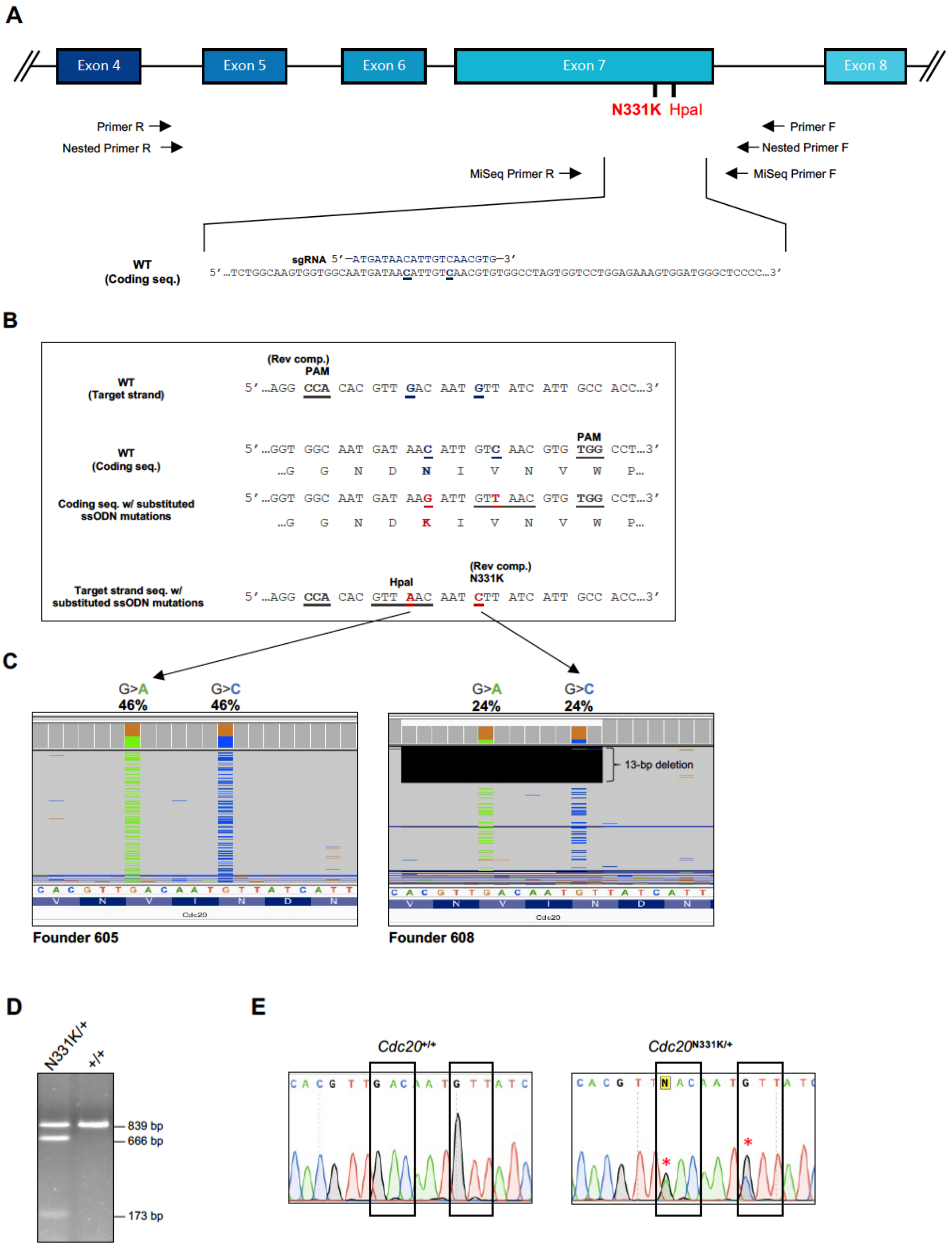


**Figure S5: Strategy for the generation of *Cdc20*$^{N331K/+}$ mice.**
**(A)** Schematic showing the CRISPR targeting strategy used to introduce the c.993G>C (N331K) mutation in the *Cdc20* locus on chromosome 4 of *Mus musculus*. The GRCm38.p4 assembly from the NCBI database was used to locate the N331 site in exon 7. The coding sequence of WT *Cdc20* is shown with the relative position of the single-stranded guide RNA (sgRNA) sequence

used for targeting. The bolded blue letters on the WT coding sequence denote the N331K and HpaI mutations sites. The relative location of the primers used for sequencing and genotyping are shown (since the *Cdc20* gene is on the reverse DNA strand in the mouse genome, the directions of the primers are shown accordingly).

**(B)** A detailed summary of the base substitutions generated using a single stranded deoxy-oligonucleotide (ssODN) repair template. The coding (reverse) strand sequence is shown with the relevant mutations and the corresponding amino acid changes. The creation of the HpaI site from a silent mutation is indicated by the grey underline in both the coding and target strand sequences.

**(C)** Illumina MiSeq results of founder C57BL/6 mice ('605' and '608') with successful incorporation of the N331K and HpaI mutations. The base substitution percentages are shown for each founder.

**(D)** Genotyping of backcrossed mice using restriction fragment length polymorphism (RFLP) analysis on HpaI-treated PCR amplicons, comparing the *Cdc20*$^{N331K/+}$ and *Cdc20*$^{+/+}$ genotypes.

**(E)** Sanger sequencing of *Cdc20*$^{N331K/+}$ and *Cdc20*$^{+/+}$ to validate our genotyping strategy. The heterozygote peaks are indicated by the red asterisks on the *Cdc20*$^{N331K/+}$ chromatogram.

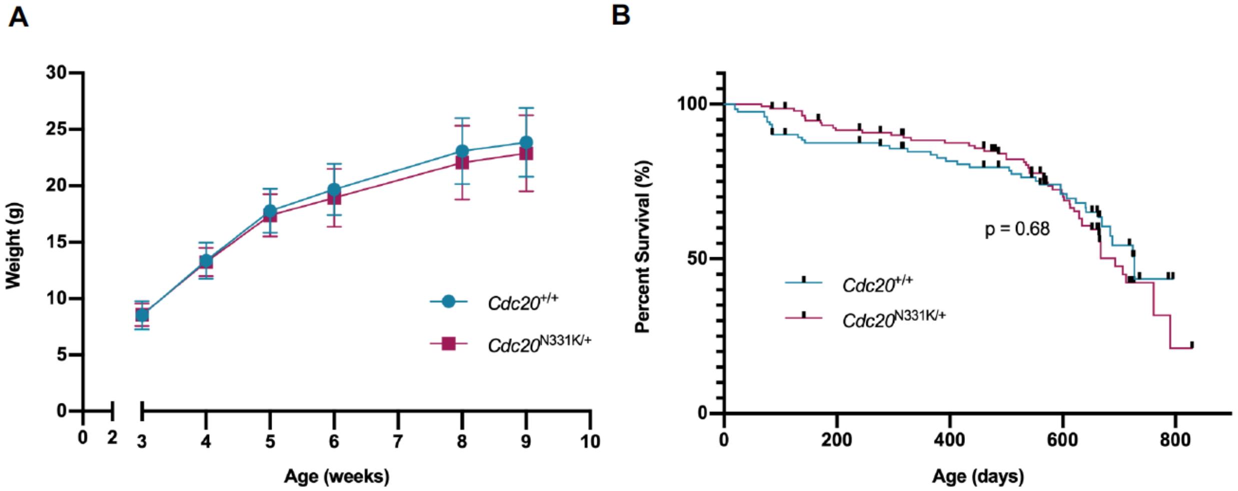


C

| Age | *Cdc20+/+* | *Cdc20N331K/+* | *Cdc20N331K/N331K* |
|---|---|---|---|
| Morula (E3.5) | 3 | 5 | 1 |
| Blastocyst (E4.5) | 3 | 5 | 8 |
| E10.5 | 2 | 11 | **0** |
| E11.5 | 3 | 3 | **0** |

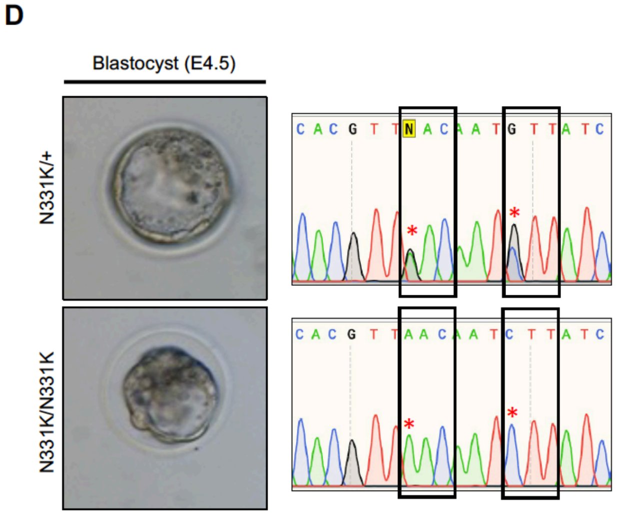


**Figure S6: Development and survival of *Cdc20*$^{N331K/+}$ mice are comparable to those of WT mice, but *Cdc20*$^{N331K/N331K}$ animals die during embryonic development.**

**(A)** Growth curves of *Cdc20*$^{N331K/+}$ and *Cdc20*$^{+/+}$ mice showing weight gained by animals during the first nine weeks after birth. *Cdc20*$^{N331K/+}$: n = 25, *Cdc20*$^{+/+}$: n = 18. Error bars: SD.

**(B)** Survival (Kaplan-Meier) plot comparing overall lifespans of WT and heterozygote animals. All mice were euthanized at clinical endpoint (inactive, hunched, presence of tumors, etc.). *Cdc20*$^{N331K/+}$: n = 145, median survival = 727 d; *Cdc20*$^{+/+}$: n = 122, median survival = 693 d; p = 0.68. Error bars = SD.

**(C)** Table showing the number of viable embryos by genotype at different timepoints during development from mating heterozygote animals. Fertilized zygotes were harvested from heterozygote females mated with heterozygote males and were cultured *in vitro* up to the blastocyst stage.

**(D)** A comparison between heterozygote and homozygote embryo morphology at the blastocyst stage. Sanger sequencing was performed to confirm the genotypes of the embryos.

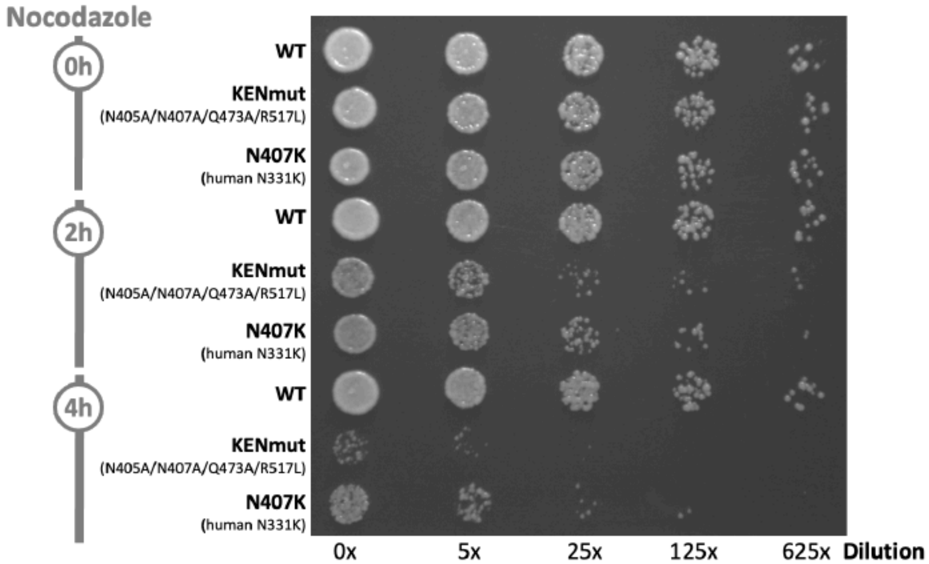


**Figure S7: The N331K variant equivalent in *S. cerevisiae* compromises the SAC and is lethal in a haploid context in a checkpoint cell survival assay.**

Endogenous WT Cdc20 in *S. cerevisiae* was replaced by a transformed duplicate WT Cdc20 copy expressed from the URA3 plasmid using the KanMX system. A LEU selection marker plasmid carrying either the N331K variant equivalent (N407K) or the KENmut (N405A/N407A/Q473A/R517L) was then transformed into the yeast and the duplicate WT Cdc20 was driven out by 5FOA treatment. For the cell survival assay, different dilutions of yeast were treated with 50 μg/ml nocodazole for up to 4 h. Compared to the WT, there was a reduction in viability of the mutants after additional of nocodazole as mutant cells kept dividing despite microtubule-kinetochore destabilization. This presumably causes genomic instability and catastrophe in progeny cells. The N407K mutant had a similar impact in disabling the SAC as mutating all four residues in the KENmut. A similar assay could not be performed for the L151 variant due to the lack of conservation between human and yeast for this residue.
